# Ketogenic diet as a metabolic vehicle for enhancing the therapeutic efficacy of mebendazole and devimistat in preclinical high-grade gliomas grown in juvenile mice

**DOI:** 10.1101/2023.06.09.544252

**Authors:** Purna Mukherjee, Jack Maurer, Sylwia A. Stopka, Bennett Greenwood, Juan J. Aristizabal-Henao, Srada Karmacharya, Alexandra Chimento, Nathan Ta, Derek C. Lee, Tomas Duraj, Roderick T. Bronson, Michael A. Kiebish, Thomas N. Seyfried

## Abstract

Invasion of high-grade glioma (HGG) cells through the brain and spinal cord is a leading cause of cancer death in children. Despite advances in treatment, survivors often suffer from lifelong adverse effects of the current toxic therapies used for management. This study investigated the influence of nutritional ketosis on the therapeutic action of mebendazole (MBZ) and devimistat (CPI-613) against the highly invasive VM-M3 and non-invasive CT-2A glioblastoma cells grown orthotopically in juvenile syngeneic mice. Additionally, both drugs were tested in the human pediatric GBM cell line SF-188. DON (6-Diazo-5-oxo-L-norleucine) was used as a positive drug control for glutamine targeting. Cerebral implantation of the VM-M3 cells, which are mesenchymal origin, invaded throughout the brain and the spinal column similar to that seen in children with HGG. Neither the CT-2A nor the VM-NM1 glioblastoma stem cell tumors showed distal invasion in syngeneic juvenile mouse brains. The maximum therapeutic benefit of MBZ and CPI-613 on tumor invasion, growth, and mouse survival occurred only when the drugs were administered together with a ketogenic diet (KD). MBZ treatment inhibited both the glutaminolysis and the glycolysis pathways in VM-M3 cells grown either *in vivo* or *in vitro*. Both MBZ and CPI-613 significantly reduced the *in vitro* growth and viability of the SF-188 cells. Moreover, drug administration together with the KD allowed for lower dosing thus minimizing toxicity while improving overall survival of the mice. This preclinical study in two different HGGs, grown in syngeneic juvenile mice, highlights the potential importance of diet/drug therapeutic strategies for managing childhood brain cancer.

## Introduction

The rise in incidence of glioblastoma (GBM) across all age groups over the last 21 years requires urgent action for management and prevention [1–3]. Diffusely infiltrating pediatric high-grade gliomas (HGGs) are the leading cause of cancer death in children in the US with no improvement in management in over 10 years [4–7]. Most survivors experience multiple adverse effects that significantly reduce quality of life [8, 9]. HGGs are classified as anaplastic astrocytoma, diffuse intrinsic pontine glioma (DIPG), and pediatric glioblastoma [10, 11]. Interestingly, glioma cell dissemination through cerebrospinal fluid (CSF) is more common in pediatric HGG patients than in adult glioma patients with similar histopathology. Distal tumor cell invasion through CSF could explain in large part the poor survival prognosis in pediatric HGG patients [6, 12].

The incidence of spinal cord spread has been reported between 2–14 % of adult glioma patients at the time of tumor recurrence [12, 13]. The pattern of recurrence of pediatric malignant gliomas can be different from that reported for adults, given the significant incidence of distant relapse and high risk of leptomeningeal dissemination [14]. Although the role of macrophage/microglia in invasion and metastasis is still an active area of research, substantial evidence shows that cells expressing microglia/macrophage markers comprise 30–50% of the cells in malignant gliomas including the tissue of DIPG and GBM [15–17]. We previously proposed that some of the microglia/macrophages present in the glioma tissue are part of the neoplastic malignant cell population and, given their mesenchymal amoeboid properties, may be capable of invading the surrounding parenchyma giving rise to tumor growth and even metastasis [18]. In that respect, our clonal population of VM-M3 cells, which arose spontaneously in the brains of the VM/Dk mice, are highly aggressive and metastatic regardless of whether the cells are implanted orthotopically or subcutaneously in the mice [19, 20]. The rapid growth and distal brain invasion of the VM-M3 tumor cells through Scherer’s structures is also remarkably similar to those features seen in human GBM [21]. In this study, we found that invasion of VM-M3 cells from the cerebral cortex into the spinal column is similar to that seen in pediatric HGG.

The response to chemotherapy is poor in most pediatric brain cancer patients with survivors experiencing unacceptable toxicity and reduced quality of life [22, 23]. Temozolomide, the most common chemotherapy for adult glioma patients, has only modest therapeutic benefit for pediatric brain cancer patients and is unacceptably toxic [24, 25]. In contrast to the complex genetic and cellular heterogeneity seen in gliomas, aerobic glucose fermentation (Warburg effect) is the common metabolic pathology seen in most if not all adult and pediatric gliomas [26–28]. In addition to aerobic glucose fermentation, brain tumor cells also depend on glutamine fermentation for growth and survival [28–30]. The reliance on glucose and glutamine for glioma malignancy comes from the well-documented defects found in mitochondrial OxPhos [31]. Glucose and glutamine fermentation in the cytoplasm and mitochondria, respectively, can compensate for OxPhos insufficiency [29, 32]. We recently showed how the glutamine antagonist, 6-diazo-5-oxo-L-norleucine (DON), administered together with a calorically restricted ketogenic diet (KD-R) could manage late-stage orthotopic growth in two adult syngeneic mouse models of GBM [33]. DON targets glutaminolysis, while the KD-R reduces glycolysis and elevates ketone bodies that are non-fermentable and neuroprotective. The diet/drug therapeutic strategy induced tumor cell death while reversing disease symptoms and improving overall survival. Moreover, the KD-R diet facilitated DON delivery to the brain thus allowing a lower dosage to achieve therapeutic effect. As drug delivery to the pediatric brain is considered a major limitation to effective HGG managements, procedures that facilitate non-toxic drug delivery to brain will have immediate and significant impact for children with brain cancer.

The antiparasitic drug, mebendazole (MBZ), showed preclinical efficacy in models of GBM and medulloblastoma [34, 35]. A phase I clinical trial for newly diagnosed HGG pediatric patients has been initiated for MBZ (ClinicalTrials.gov Identifier: NCT01837862). The aim was to evaluate safety and dosing in treating pediatric patients with recurrent or refractory gliomas. MBZ treatment *in vivo* as a single agent or in combination with chemotherapy led to the reduction or complete arrest of tumor growth, marked decrease in metastatic spread, and increase in overall survival [36–38]. The mechanism by which MBZ inhibits tumor growth is due in part to the inhibition of glycolytic enzymes and glucose uptake [39, 40]. MBZ has relatively low toxicity in both adults and in children [41–43]. The systemic bioavailability of MBZ is enhanced when co-administered with fatty foods [44, 45]. In this study, we administered MBZ with a high-fat, low carbohydrate ketogenic diet to juvenile mice implanted orthotopically with the VM-M3 and CT-2A malignant gliomas. The lipoate derivative CPI-613 is a novel anti-cancer agent that targets mitochondrial dehydrogenase activities (pyruvate dehydrogenase and α-ketoglutarate dehydrogenase/2-oxoglutarate dehydrogenase, OGDH) causing apoptosis, necrosis, and autophagy of tumor cells [46–50]. CPI-613 had little toxicity in therapeutic dose ranges and was well-tolerated at higher doses in preclinical models (the maximum tolerated dose in mice was 100 mg/kg). CPI-613 (10 mg/kg) significantly inhibited the growth of the H460 human non-small cell lung carcinoma in xenograft mice. CPI-613 also caused robust tumor growth inhibition in a mouse model of human pancreatic tumor (BxPC-3) [51]. Although CPI-613 was therapeutic against preclinical cancers, it failed to show non-toxic therapeutic efficacy against metastatic pancreatic cancer and relapsed or refractory acute myeloid leukemia in two phase III clinical trials [52–54]. The influence of CPI-613 on TCA cycle metabolites was recently studied in preclinical model of brain tumors [55]. However, there are no reports to our knowledge that have evaluated an effect of CPI-613, used alone or in combination with other drugs or diets, on pediatric HGG.

Here we show for the first time that MBZ and CPI-613 are each therapeutically synergistic when administered together with a ketogenic diet in juvenile mice in the invasive VM-M3 and in the non-invasive CT-2A stem cell glioma models. Moreover, we report that MBZ can target the glutaminolysis pathway of VM-M3 cells both *in vitro* and *in vivo*. These findings further support the dependency of the invasive VM-M3 glioma cells on glutamine for growth. Both drugs prevented dissemination of the VM-M3 tumor cells from the brain to the spinal cord when administered with the KD in juvenile mice. Moreover, lower drug dosages could achieve remarkable non-toxic improvement in overall survival when administered with the KD. This novel diet/drug cocktail therapy could have significant translational impact for children with malignant brain cancers. An initial report of these findings was presented [56].

## Results

The aim of this research was to determine if a ketogenic diet could improve the therapeutic efficacy of MBZ and CPI-613 in two preclinical glioblastoma models that were grown orthotopically in juvenile mice (postnatal day 21-25). The mouse brain at these ages is comparable to that of children between 3-7 years of age [57]. None of the mice used in this study received radiation or steroids as part of the metabolic treatment protocol to avoid the provocative effects of elevated blood glucose, immune-suppression, and systemic inflammation on tumor growth [58, 59]. *In vivo* bioluminescence imaging, reflecting tumor progression, was measured as a primary end point while overall mouse survival was measured as a secondary end point. DON treatment alone was used as a positive control throughout the study based on our previous observations [33]. The juvenile mice were fed *ad libitum* KD to maintain body weight. The models included the highly invasive VM-M3 tumor of mesenchymal origin and the highly angiogenic, but non-invasive, CT-2A tumor of neural stem cell origin grown in their respective VM/Dk and C57BL/6J inbred syngeneic hosts [20]. The VM-NM1 tumor, which is similar in cell origin and angiogenesis phenotype to the CT-2A tumor, was also evaluated. The timeline of the study from tumor implantation in the brain to the initiation of diet/drug administration through termination or survival is shown in **Figure 1**. The diet/drug treatment protocol used for this study was based on the Press-Pulse therapeutic strategy that we described previously [60].

**Figure 1.**
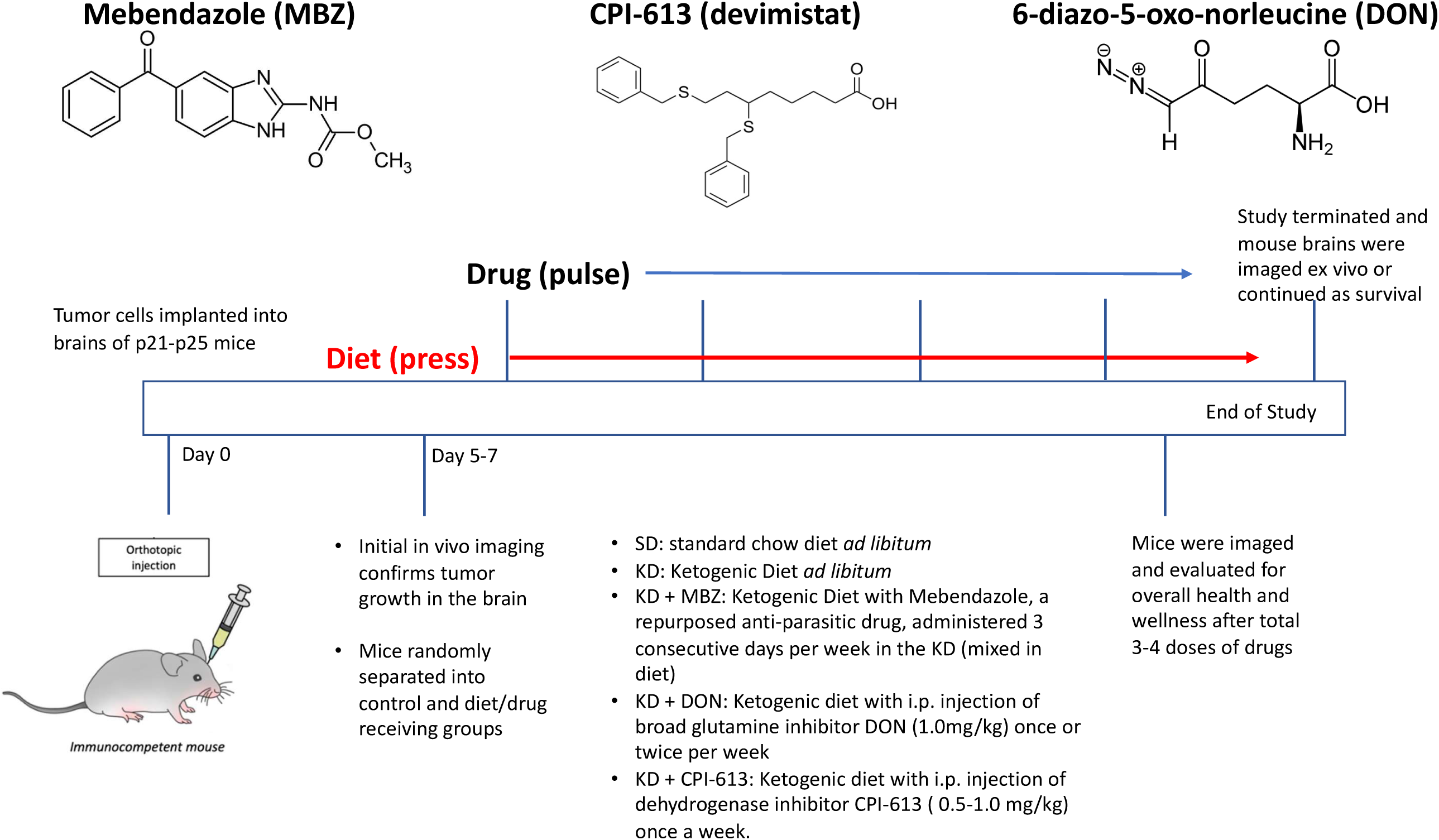
Experimental design for the analysis of diet/drug influence on brain tumor growth and survival in juvenile mice. Juvenile mice (p21-p25) were used as hosts for the orthotopic implantation of the highly invasive VM/M3 and the non-invasive CT-2A brain tumors as detailed in the text. A ketogenic diet (KD) was employed as a press therapy throughout the study as previously described for press-pulse therapy [60]. The KD maintains a lower blood glucose/ketone index (GKI) compared to the high carbohydrate control diet [19]. Mebendazole (MBZ), CPI-163 (devimistat), and DON were used as a pulse therapy and were dosed and scheduled according to maintain maximum mouse health. *In vivo* bioluminescence imaging of the mice was as described in methods. *Ex vivo* bioluminescence imaging and brain collection was done on the termination day. For the survival study, the same protocol was followed, and individual mouse deaths were recorded as described in the text. The experiments were repeated when mouse litters became available for study. The data obtained from each experiment were then presented individually or pooled for statistical analysis.

### VM-M3 glioma cells invade the spinal cord from the cerebrum implantation site in juvenile mice

VM-M3 tumor cells (100,000 cells in 5.0 µL in phosphate buffered saline, PBS) were implanted into the brains of juvenile postnatal day p21-25 mice of the syngeneic VM/Dk inbred strain. To verify tumor take, mice were imaged using the luciferin-luciferase bioluminescence reporter system on day 6 after tumor implantation. It is interesting to note that the VM-M3 tumor cells invaded the spinal cord from the brain implantation site (**Figure 2A & 2B**). We have not previously seen spinal cord invasion of VM-M3 tumor cells when implanted into the cerebrum of adult VM/Dk mice [33, 61]. Furthermore, spinal cord invasion was not observed following brain implantation of the non-invasive VM-NM1 glioma stem cell tumor into juvenile mice (**Figure 2C**). It is also important to mention that cerebrospinal dissemination of malignant glioma is more common in children than in adults and is indicative of poor survival [14]. Hence, our juvenile pre-clinical VM-M3 glioma model is relevant to the clinical situation seen in children with HGG.

**Figure 2.**
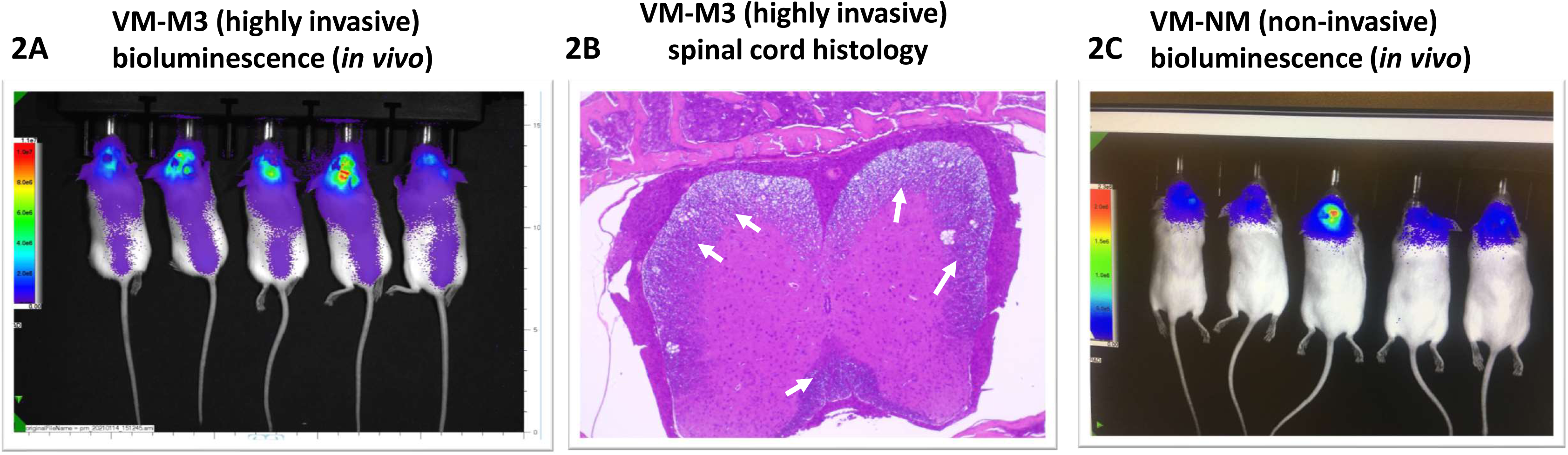
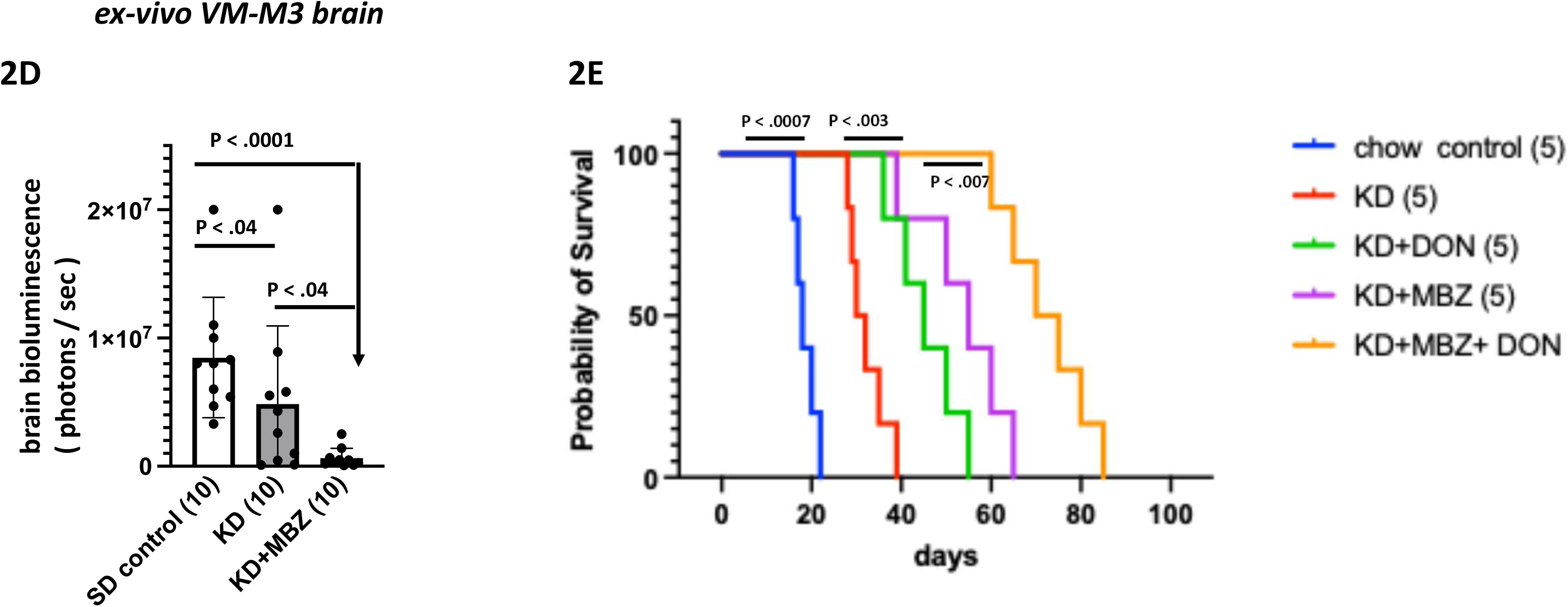
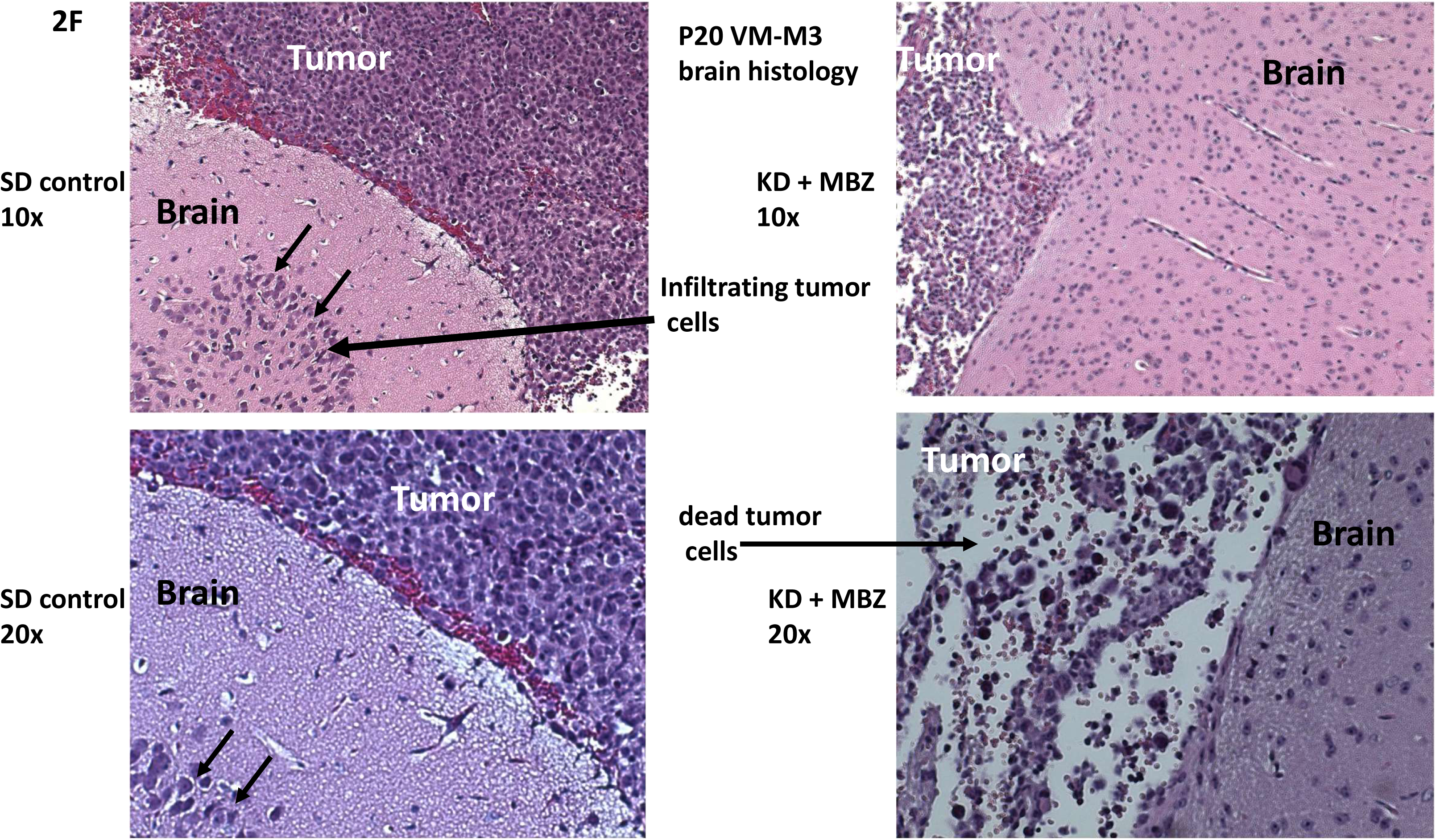
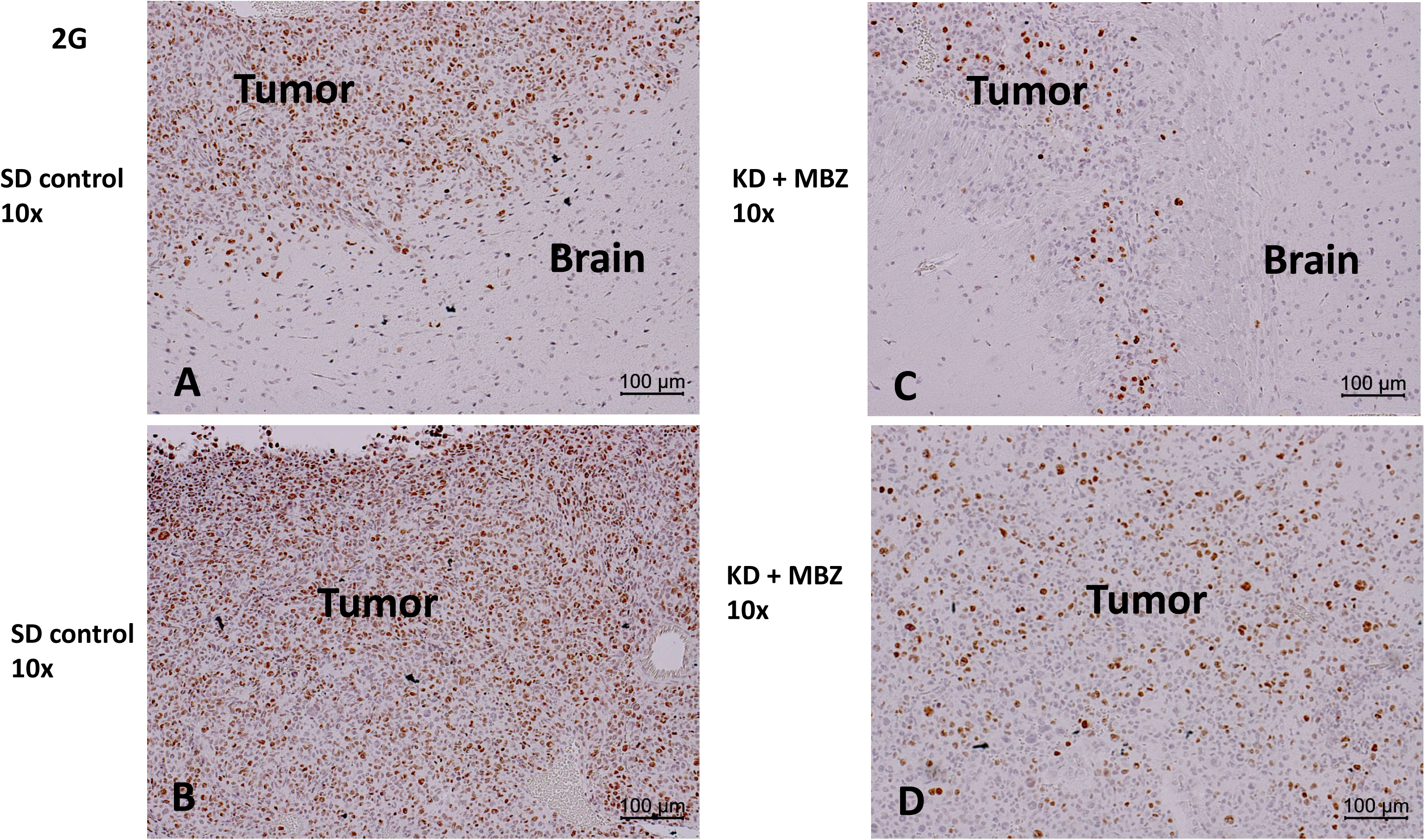
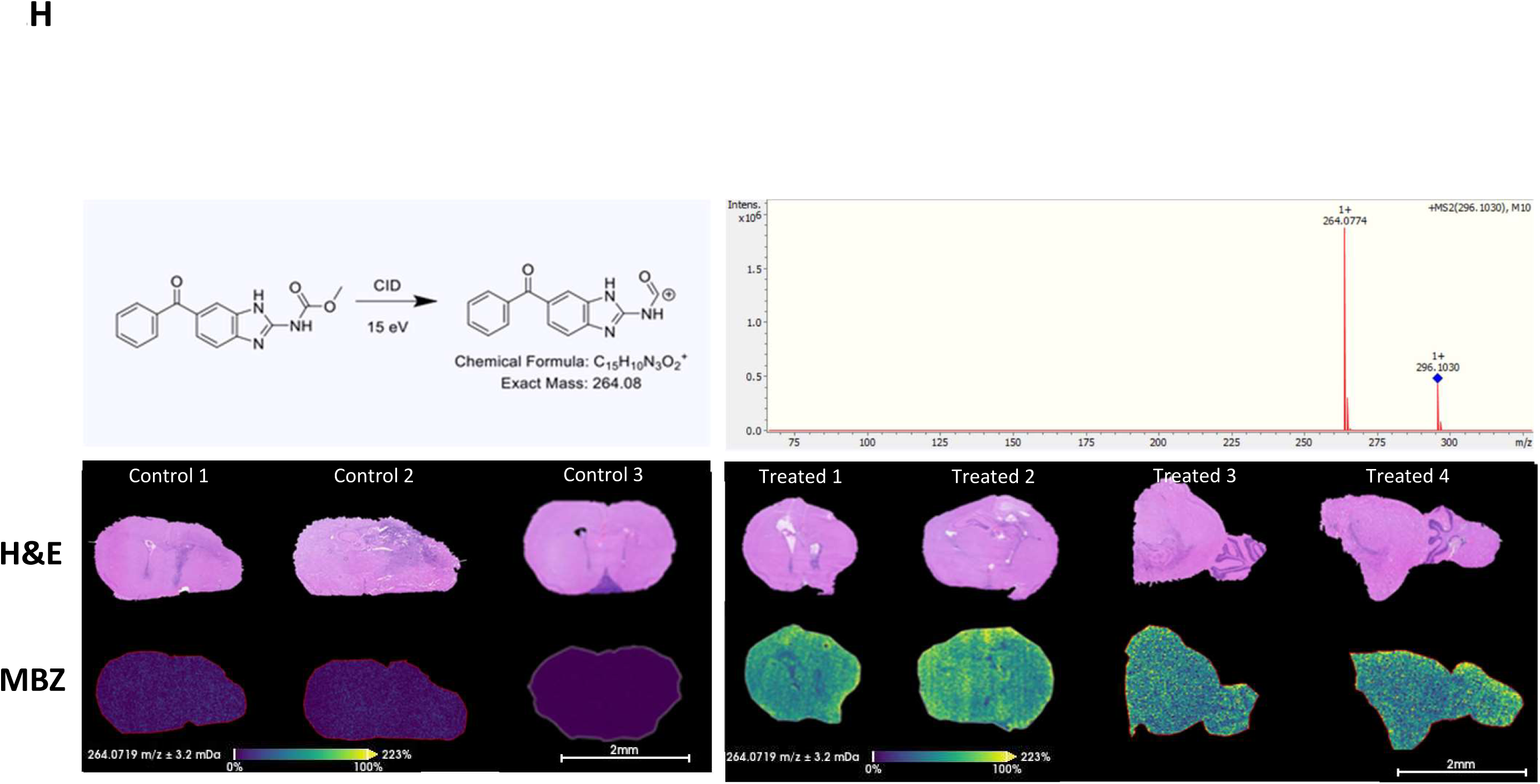
Effects of MBZ used with KD on orthotopically implanted VM-M3/luc glioblastoma cells in the brain of p20-25 VM/Dk mice. **A.** VM-M3/luc tumor cells invasion from the brain implantation site to the spinal column. This finding is indicative of tumor cell invasion through CSF (*in vivo* analysis). **B.** Histological analysis of tumor cell invasion in the spinal cord. Dense infiltrates of tumor cells are seen in the spinal cord and meninges. **C.** Brain growth of the non-invasive stem cell VM-NM/luc glioma grown in juvenile mice. Spinal cord invasion was not seen following brain implantation. **D.** Evaluation of *ex vivo* bioluminescence of brains of mice with diet and drugs. Brain bioluminescence was significantly lower in the MBZ-treated mice than in either the SD control or the KD alone mouse groups. Values are expressed as the mean ± SEM and Mann-Whitney test was used to determine the significance between the groups. **E.** The Kaplan-Meier survival data showed that overall survival was longer in the KD-fed mice than SD control mice, but that overall survival was significantly longer in mice receiving the KD containing either MBZ or DON than in mice receiving the KD alone. Log-rank (Mantel-Cox test) analysis was performed to determine the significance between groups. **F.** The p20 mice were implanted with VM-M3/luc tumor cells into the cerebral cortex and brain tissue histology (H&E) evaluated 12 days later. The low power (10x) and higher power (20x) images on the left show many VM-M3/luc cells invading into normal appearing brain tissue in a mouse fed the SD control diet. Tumor cell death without obvious invasion is shown in the low power (10x) and higher power (20x) images on the right in a mouse fed the KD + MBZ diet. **G.** Ki-67 staining shows a marked reduction in positive cells in the treated tissue at both the brain-tumor junction (**C**) and the primary tumor mass (**D**) compared to the untreated brain tumor (**A** and **B**). 3 independent mice brain tissues in each group were stained for both histology and immunohistochemistry. **H.** MALDI imaging mass spectrometry analysis revealed detectable MBZ (transition *m/*z 296→264) in the treated brains supporting effective delivery and tissue accumulation of MBZ. Same sections were also stained with H&E subsequently to MALDI imaging. 4 independent mice for MBZ treated brains and 3 untreated control brains were used for the study. 3 control brains (lower panel, left) did not show the yellow pixel that associated with the presence of MBZ. In contrast, all 4 treated brains displayed a marked increase in yellow pixels confirming the presence of the drug in the brain tissue. These findings further suggest that administration of MBZ with a KD could offer a powerful non-toxic therapy for managing HGG in children.

### The KD enhances the therapeutic effects of MBZ against the VM-M3 glioma by reducing growth in the brain and by inhibiting invasive spread to the spinal cord in juvenile mice

After confirmation of tumor take using bioluminescence *in vivo* imaging, the mice were divided into the diet/drug groups. MBZ (100 mg/kg body weight) was administered to the mice in the KD for 3 consecutive days/wk. The KD, without MBZ, was administered continuously throughout the study. The study was terminated when the mice in the standard diet-fed (SD) control group lost about 20% body weight and developed signs of morbidity. All mice were imaged to evaluate brain tumor growth *in vivo* before removal of the brains for *ex vivo* imaging. DON (1.0 mg/kg body weight), used as a positive control, was injected intraperitoneal (i.p.) once per week. Each experiment was repeated 3-4 times in the same way when mouse litters became available. Male and female mice from each litter were randomly distributed among the groups, i.e., both males and females were tested for each KD/drug combination. We chose not to evaluate the efficacy of these drugs administered in the high-carbohydrate standard mouse diet (SD) based on previous studies showing that therapeutic efficacy of MBZ and DON is improved when the drugs are administered in high-fat diets [33, 35].

A preliminary study was conducted on a single litter of juvenile mice that included a mouse standard diet (SD) control, a KD control, a KD + MBZ experimental, and a KD + DON experimental. This study involved both *in vivo* and *ex vivo* imaging of brains. The imaging data **(Extended Figure 2a**) showed that brain bioluminescence, indicated as number of photons × 10^x^ was lower both *in vivo* and *ex vivo,* in the mice treated with KD + MBZ or with KD + DON than in mice receiving he KD or the SD alone. We found that both MBZ and DON prevented tumor cell spread through the CSF when administered with KD. Bioluminescence was greater on the right ipsilateral side (tumor implanted side) than on the left contralateral side of the brains. Bioluminescence detected on the left contralateral side is indicative of distal hemispheric tumor cell invasion as previously described [21]. Results from a repeat study showed that brain bioluminescence was significantly lower in the mice receiving KD + MBZ than in the mice receiving either the SD or the KD alone suggesting a significant reduction in tumor growth in the juvenile brain after MBZ treatment (**Figure 2D)**. The survival study was done in a separate group of VM p21 mice. Overall survival was longer in the KD-fed mice than in the SD-fed control mice while the overall survival was significantly longer in mice receiving the KD containing either MBZ or DON than in mice receiving the KD alone (**Figure 2E**). Although a synergistic effect of MBZ and DON was observed on mouse survival, we chose to focus on MBZ for further experiments, as it is the main subject of the current study. In addition, DON is not an approved or readily available drug and is unlikely to be used in children at this time. The video in the supplementary material shows that the mice on KD/drug (KD + MBZ + DON) were ambulatory and healthy in contrast to the mice fed the control high-carbohydrate SD that were sedentary and moribund (**Extended Figure 2b, video file**). Histological examination of liver tissues from the KD + MBZ group showed no toxicity-related changes in tissue architecture, such as enlarged hepatocytes, necrosis, or immune cell infiltration associated with inflammation (**Extended Figure 2c**). Serum alanine transaminase activity (ALT) levels were comparable between the two groups **(Extended Figure 2d).** Histopathological analysis revealed tumor cell death without obvious invasion in the brains of a mouse fed the KD + MBZ diet (**Figure 2F**). Ki-67, a proliferation marker of tumor cells, was assessed by immunostaining in brain tumor tissues from SD-fed control mice and diet/drug-treated mice. Results showed a marked reduction in the number of Ki-67 positive cells in the treated mice compared with control mice. This reduction was evident in both the brain–tumor junction and the implantation site where the primary tumor mass was present (Figure 2G). No statistically significant differences in the therapeutic benefits of diet drug cocktails on tumors between male and female mice.

### MBZ detection in VM/M3 Brain Tumor Tissue by MALDI Mass Spectrometry and Microscopy Imaging

A separate study was conducted to detect MBZ in brain tissue. KD-fed mice were administered MBZ using the same dose and schedule previously described in Figure 2. After termination, the brains were flash-frozen for imaging purposes as described [62]. MALDI imaging mass spectrometry analysis revealed detectable levels of MBZ (transition m/z 296→264) in the MBZ treated brains. Untreated SD brains were processed and imaged as negative controls. These findings support the effective delivery and accumulation of MBZ in brain tissue (**(Figure 2H)**.

### MBZ targets both the glutaminolysis and the glycolysis pathways in VM-M3 glioma cells

MBZ treatment reduced the bioluminescence in the VM-M3 cells in a dose dependent manner indicating reduced ATP content and reduced cell growth/viability (**Figure 3A**). DON, used as a positive control, also reduced the bioluminescence of VM-M3 cells (**Figure 3B**). In contrast to MBZ, which showed a dosage effect for viability, no dosage effect for proliferation was seen for DON. Based on this information, we used 1.0 μM of MBZ and 50.0 μM of DON in further experiments. Bioluminescence was significantly lower for both MBZ- and DON-treated VM-M3 cells after 24 hours (**Figure 3C**). Brightfield images showed MBZ induced cell death in the VM-M3 cells at 10 μM whereas DON reduced the proliferation rate *in vitro* (**Extended Figure 3a**).

**Figure 3.**
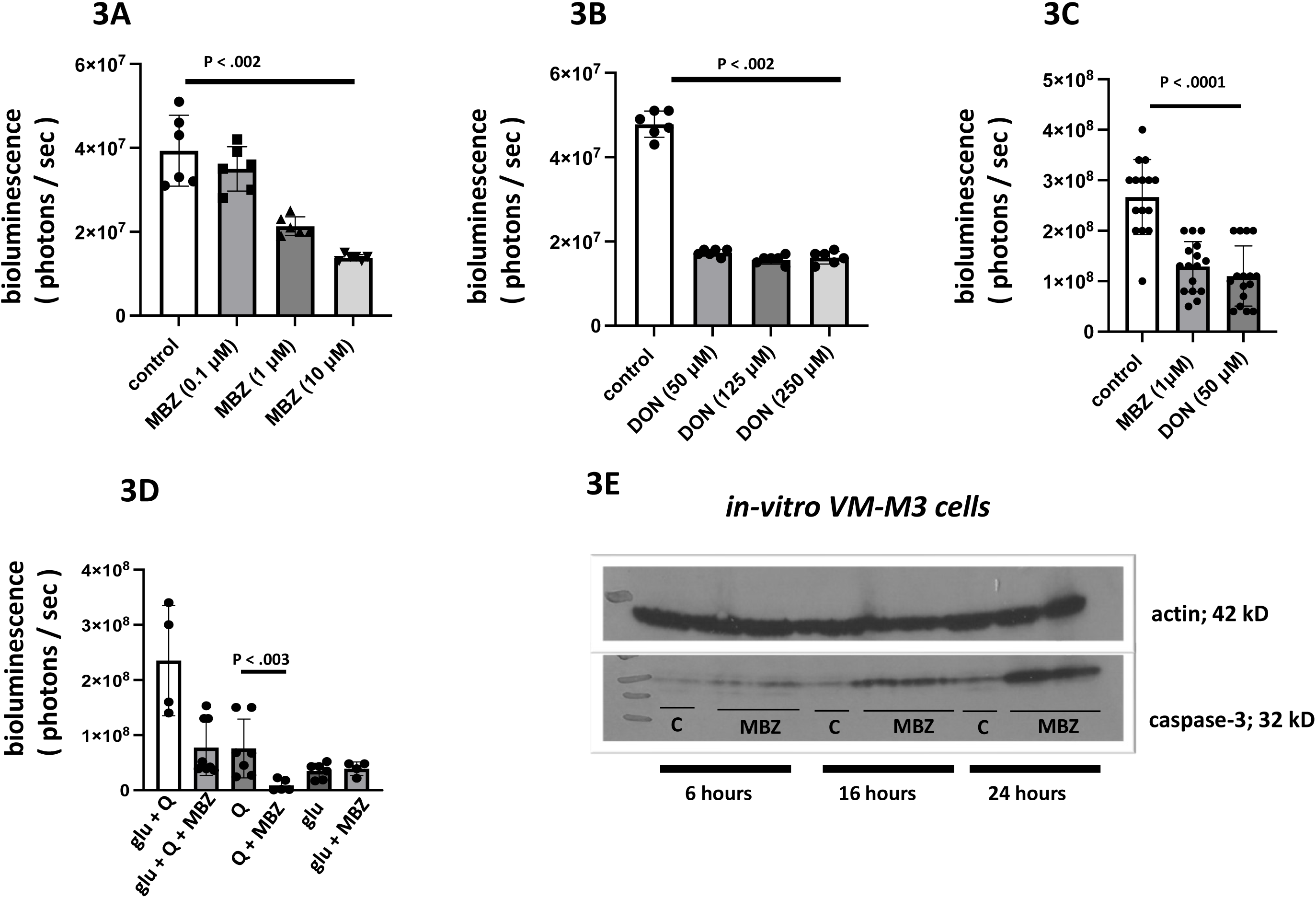
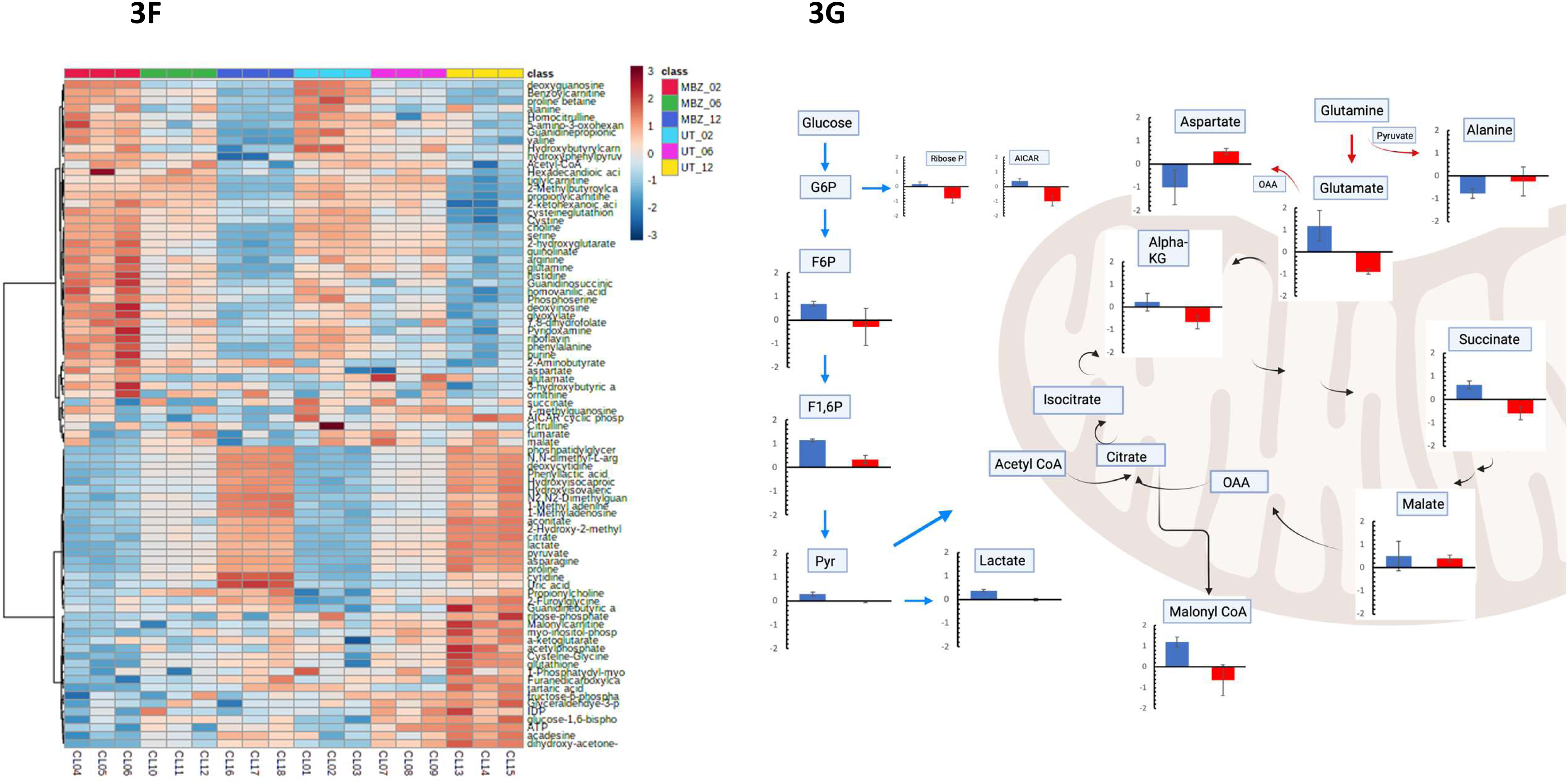
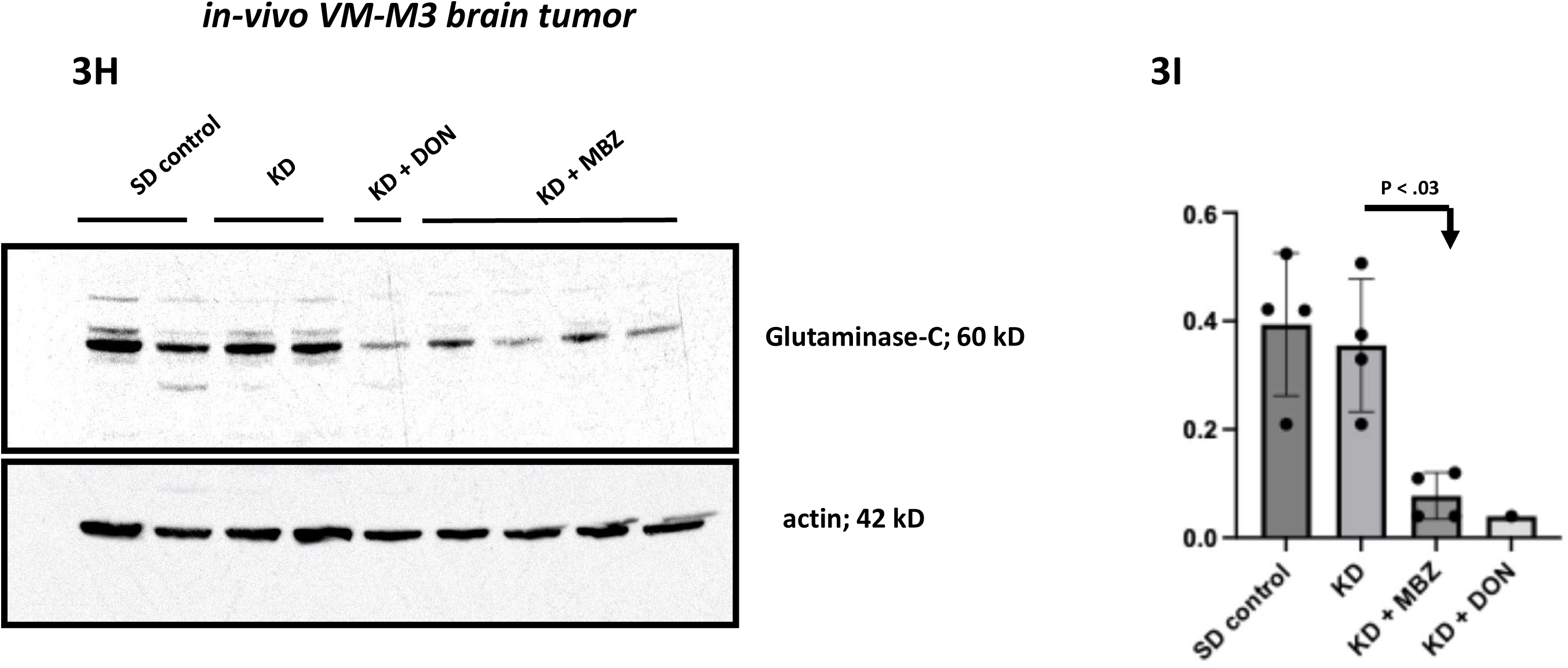
MBZ targets both the glutaminolysis and the glycolysis pathways in cultured VM-M3/luc cells. VM-M3/luc cells were treated with different doses of MBZ and DON while grown *in vitro*. DON was used as a positive control. **A-C.** MBZ reduced the bioluminescence in the VM-M3/luc cells in a dose dependent manner. Both MBZ and DON reduced the bioluminescence significantly compared to controls. Experiments were repeated three times and values are expressed as the mean ± SEM and Mann-Whitney test to determine the significance between the groups. **D.** VM-M3/luc cells are more glutamine dependent than glucose dependent for proliferation. Moreover, MBZ reduced the bioluminescence when glutamine was the only substrate in the media as explained in the results. **E.** MBZ (1μM) induced apoptosis in VM-M3 cells in a time dependent manner indicated by caspase-3 activation. **F-G.** The LC-MS metabolite data from the cell lysates showed that MBZ reduced the glutaminolysis and the glycolysis pathways. Heat map analysis revealed that levels of glutamate, α-ketoglutarate, succinate, and lactate were lower in the MBZ treated cells (red bars) than in the untreated cells (blue bars). The data are presented as normalized concentrations from three independent metabolite samples collected after 6 hours of treatment. **H-I.** Mouse brains analyzed for glutaminase C expression. Brain glutaminase C expression was significantly lower in the MBZ treated mice than in the untreated mice. DON was used as a positive control for these experiments.

To study the effect of MBZ on the glycolysis and the glutaminolysis pathways, we evaluated the effect of MBZ on VM-M3 cell proliferation in serum free basal media with and without added glucose or glutamine. VM-M3 cells (1.0 × 10^5^) were seeded in 24-well plates in complete medium as before. After 24 hours, the cells were rinsed with PBS and treated with MBZ (1.0 μM) and DON (50.0 μM) in basal media for another 24 hours. It was evident that both glutamine and glucose were required for VM-M3 proliferation (**Figure 3D**). An extensive analysis of this requirement for glioma growth was recently presented [32]. MBZ treatment significantly reduced the bioluminescence in VM-M3 cells as shown (**Figure 3C**). Interestingly, the inhibitory effect of MBZ was evident when glutamine was the main fuel present in the media, but not when glucose was the main fuel **(Figure 3D**). These findings suggest that MBZ is more effective in inhibiting glutamine-linked viability than in inhibiting glucose-linked viability of the VM-M3 cells and are consistent with previous findings with these cells [32]. Further studies will be needed to determine how MBZ might target the glutaminolysis pathway in the VM-M3 glioma cells. MBZ-treated (1.0 μM) cells showed an increase in caspase-3 expression in a time-dependent manner, suggesting that cell death is likely due to activation of apoptotic signaling pathways **(Figure 3E**). In the next experiment, VM-M3 cells were seeded in 6-well plates. After 24 hours, media was replaced and the cells were treated with MBZ (1.0 μM) for 2-, 6-, and 12-hours in serum free basal media. The LC-MS metabolite data showed that lactate levels and intermediates of glycolysis and nucleotide biosynthesis were lower in the MBZ treated cells than in the untreated cells. Glutamate and α-ketoglutarate, metabolites in the glutaminolysis pathway, were also lower in the MBZ treated cells than in the untreated cells after 6 hours. Consistent with lower levels of TCA cycle intermediates, MBZ treated cells exhibited a lower level of fatty acid biosynthesis, i.e., malonyl-CoA. The levels of 2-aminobutyrate and aspartate were higher in the MBZ-treated cells than in the untreated cells (**Figures 3F & 3G**). These findings further suggest that glutamine usage is important for the growth of VM-M3 tumor cells. Restricted glutamine usage could also account in part for the lower levels seen for succinate, the end-product of the glutaminolysis pathway [29, 31, 32]. MBZ caused a reduction in glutaminolysis metabolites. This reduction could reduce ATP production through mitochondrial substrate level phosphorylation at the succinyl-CoA ligase step in the TCA cycle as previously described [29, 31, 32, 63]. A reduction in succinate would reduce the stabilization of hypoxia-inducible factor 1α (Hif-1α) [31], which could reduce metabolite levels in the glycolytic pathway, as reflected at 6 hours (**Figures 3F & 3G**). Glutaminase C is the gatekeeper of the glutaminolysis pathway that converts glutamine to glutamate and is overexpressed in aggressive brain tumors compared to normal brains [64]. We found that Glutaminase C is overexpressed in VM-M3 and CT-2A brain tumors compared to the normal VM and B6 mouse brain tissues (**Extended Figure 3b)**. Interestingly, Glutaminase C expression was significantly lower in the MBZ treated brain tumor than in the control untreated tumor (**Figure 3H and 3I**). DON treated brain was used as a positive control which is known as a non-selective inhibitor of the enzyme [65]. These results suggest that MBZ may target both the glutaminolysis and the glycolysis pathways in the VM-M3 tumor cells.

### CPI-613 reduces the growth of VM-M3 cells *in vitro* and *in vivo*

VM-M3 cells were treated with two different doses (100 μM or 250 μM) of CPI-613. CPI-613 reduced the bioluminescence of the VM-M3 tumor cells in a dose dependent manner (**Figure 4A**). Moreover, CPI-613 (100 μM) reduced the bioluminescence by approximately 50% after 24 hours. VM-M3 cells (1.0 × 10^5^ cells in 5 µL) were implanted into the brains of juvenile postnatal-day 20-25 mice of the inbred VM/Dk strain, as described above. The mice were imaged on day 6 after tumor implantation. After bioluminescence imaging, the mice were divided into four groups that included: A SD control; SD + CPI-613; KD; and KD + CPI-613 (0.5-1.0 mg/kg body weight in DMSO) injected intraperitoneal (i.p.) once/wk. Both the SD controls and the KD- fed mice received injection of DMSO as vehicle. The mice were imaged *in vivo* on day 13 after tumor implantation. Bioluminescence was robust in mice fed the SD and in mice fed the SD + CPI-613 indicating that CPI-613 did not inhibit VM-M3 cell growth *in vivo* (**Extended Figure 4a**). On termination, *ex vivo* bioluminescence measurements were lower in the mice fed KD + CPI than in the other groups suggesting a synergistic therapeutic effect of CPI-613 with the KD (**Figure 4B**). These findings are important, as no prior studies have evaluated the therapeutic action of CPI-613 against brain cancer when administered with the KD. OGDH (2-oxoglutarate dehydrogenase) is one of three enzymes in the α-ketoglutarate dehydrogenase complex responsible for catalyzing a rate-regulating step of the TCA cycle. CPI-613 is known to reduce OGDH protein expression in pancreatic cancer (PDAC) cells [66]. We also found that OGDH protein expression was significantly reduced in CPI-613 treated brains of KD mice (**Figure 4C and 4D**). This result suggests that the KD might facilitate the delivery of CPI-613 through the blood-brain barrier as we described previously for other small molecules [33, 67]. LC-MS MALDI imaging analysis of the brain could not detect the drug due to its water-insoluble nature. However, a significant amount of the drug was detected in the blood of mice 45 minutes after injection (Extended Figure 4b).

**Figure 4.**
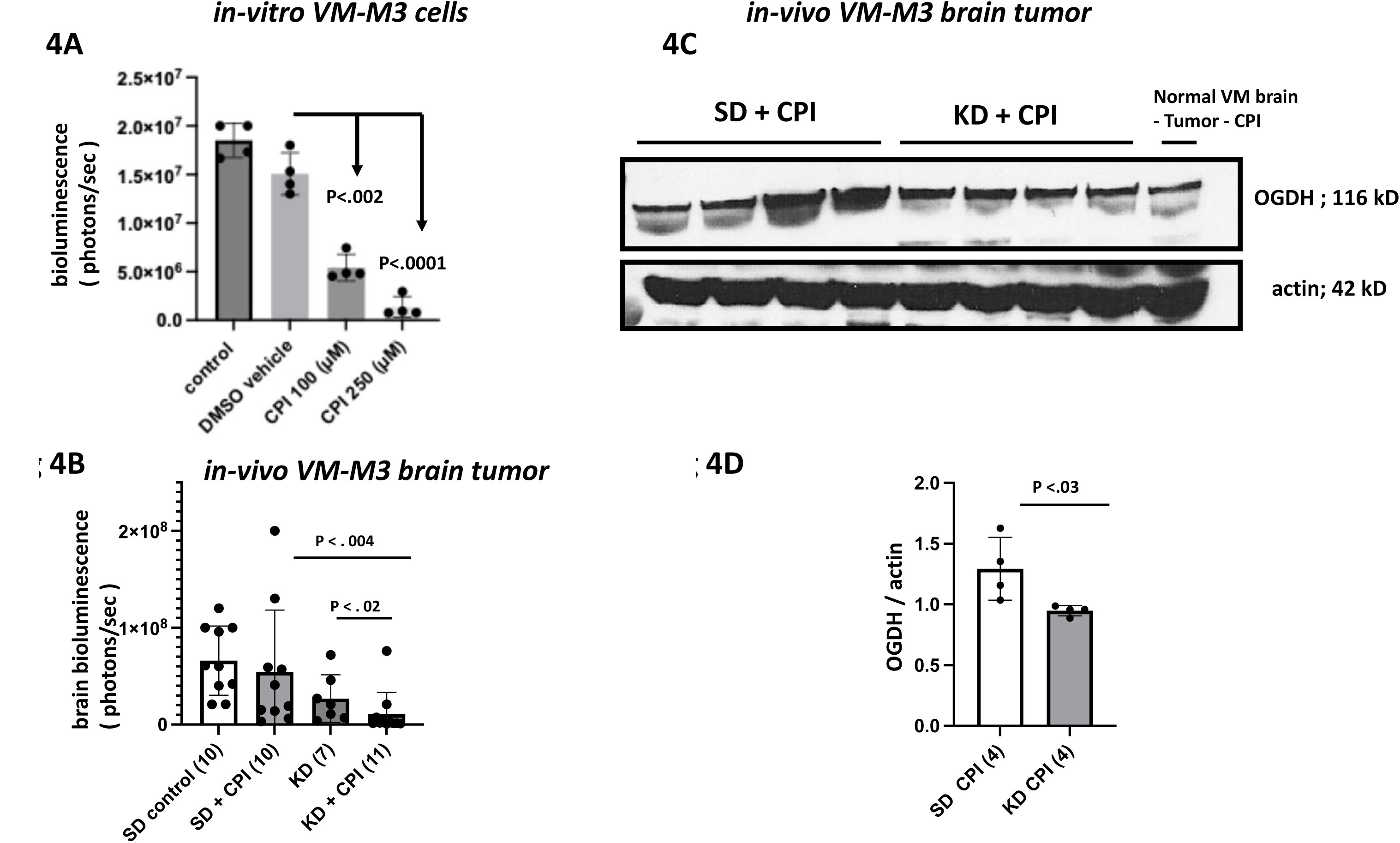
Effects of CPI-613 (devimistat) on the progression of VM-M3/luc cells *in vitro* and *in vivo.* VM-M3/luc cells were treated with different doses of CPI-613 while grown *in vitro*. **A**. CPI-613 reduced the bioluminescence in the VM-M3/luc cells in a dose dependent manner**. B.** *Ex vivo* brain bioluminescence was lowest in KD + CPI group. Values are expressed as the mean ± SEM and Mann-Whitney test to determine the significance between the groups. **C-D**. Brain OGDH expression was significantly lower in the KD + CPI-613 treated mice than in the than in the brains of mice SD + 613.

In addition to reduction in bioluminescence which links directly to ATP synthesis, MBZ and CPI-613 significantly reduced oxygen consumption rate (OCR). To determine the OCR, cells were treated with MBZ (1μM) and CPI-613 (100μM) for 6 hours in DMEM containing 12 mM glucose and 2 mM glutamine without serum. A significant reduction in OCR was recorded in real time from both drugs treatment at 6 hours compared to untreated control cells. (Extended Figure 4c).This demonstrates that both MBZ and CPI have direct effect on the metabolic activity of cancer cells .

### MBZ and CPI-613 enhances the survival of juvenile mice with orthotopic CT-2A tumor while under KD

In addition to evaluating the therapeutic effects of MBZ and CPI-613 on the orthotopic growth of the VM-M3 glioblastoma cells in syngeneic p21-p25 mice, we also evaluated their effect on the orthotopic growth of the CT-2A stem cell glioma in the syngeneic C57BL/6J host strain at p21-25. Overall mouse survival was significantly longer when MBZ was used together with the KD than diet alone group (**Figure 5A**). Considered together, these results indicate that the therapeutic action of MBZ is significantly improved when administered together with a KD for managing the *in vivo* growth of either the VM-M3 or CT-2A glioma cells in juvenile mice. CPI-613 treated mice also showed a significant increase in survival compared to the mice in the SD or KD control groups (**Figure 5B**). A SD drug group for the survival was not included considering that the therapeutic effect on tumor growth was better when the drug was used with KD in our highly invasive VM-M3 glioma model.

**Figure 5.**
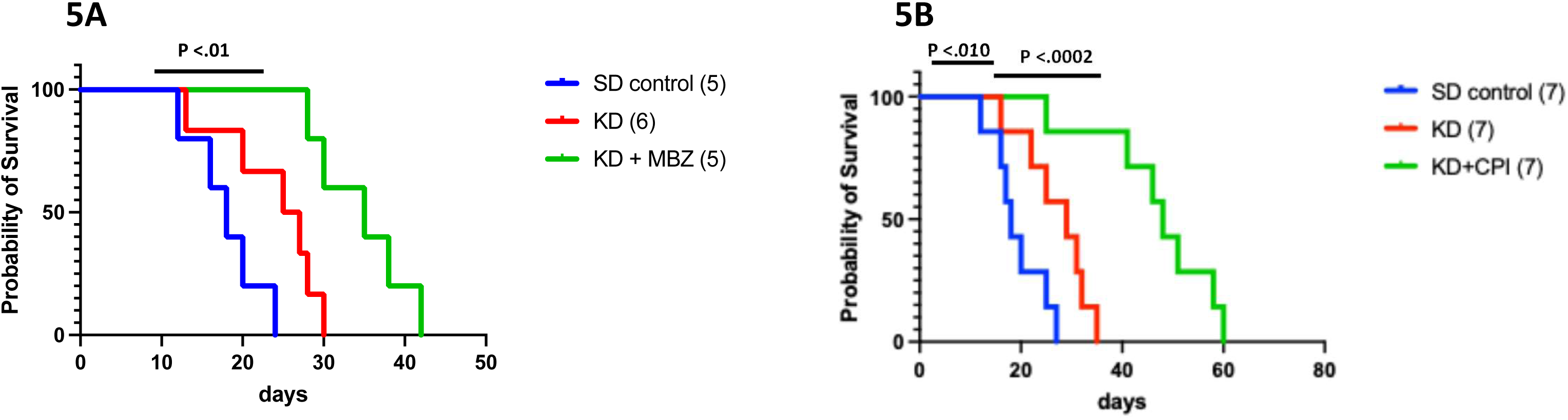
Effects of MBZ and CPI-613 (devimistat) on the survival of juvenile mice with orthotopic CT-2A tumor. CT-2A/luc glioma cells were implanted in the brains of juvenile C57BL/6 mice as described in methods. In separate survival studies, mice were treated with MBZ and CPI-613. **A.** Kaplan-Meier survival analysis revealed significantly higher survival rates in MBZ- treated and **B.** CPI-613-treated mice with CT-2A tumors in the brains compared to diet control groups. DON was utilized as a positive control. Log-rank (Mantel-Cox test) analysis was performed to determine the significance between groups.

### The KD maintains bodyweight and elevates blood β-hydroxybutyrate (β-OHB) levels while reducing blood glucose and the glucose/ketone index (GKI) in juvenile mice

Juvenile mice (p25) were tested for the influence of *ad-libitum* KD on body weight, blood glucose and β-OHB levels, and the calculated glucose ketone index (GKI) [68]. In a preliminary study, both male and female juvenile mice were fed the KD for a week to record the body weight. The body weights of the SD-fed mice increased while the body weights in the KD-fed mice remained stable (**Figure 6A & 6B**). In another set of experiments, mice were fed the KD for two weeks after which blood glucose and β-OHB levels were measured. Following these measurements, the mice were implanted with VM-M3 glioma cells as mentioned above. The diet was continued for another two weeks. The brains were removed and fixed in formalin for histology.

**Figure 6.**
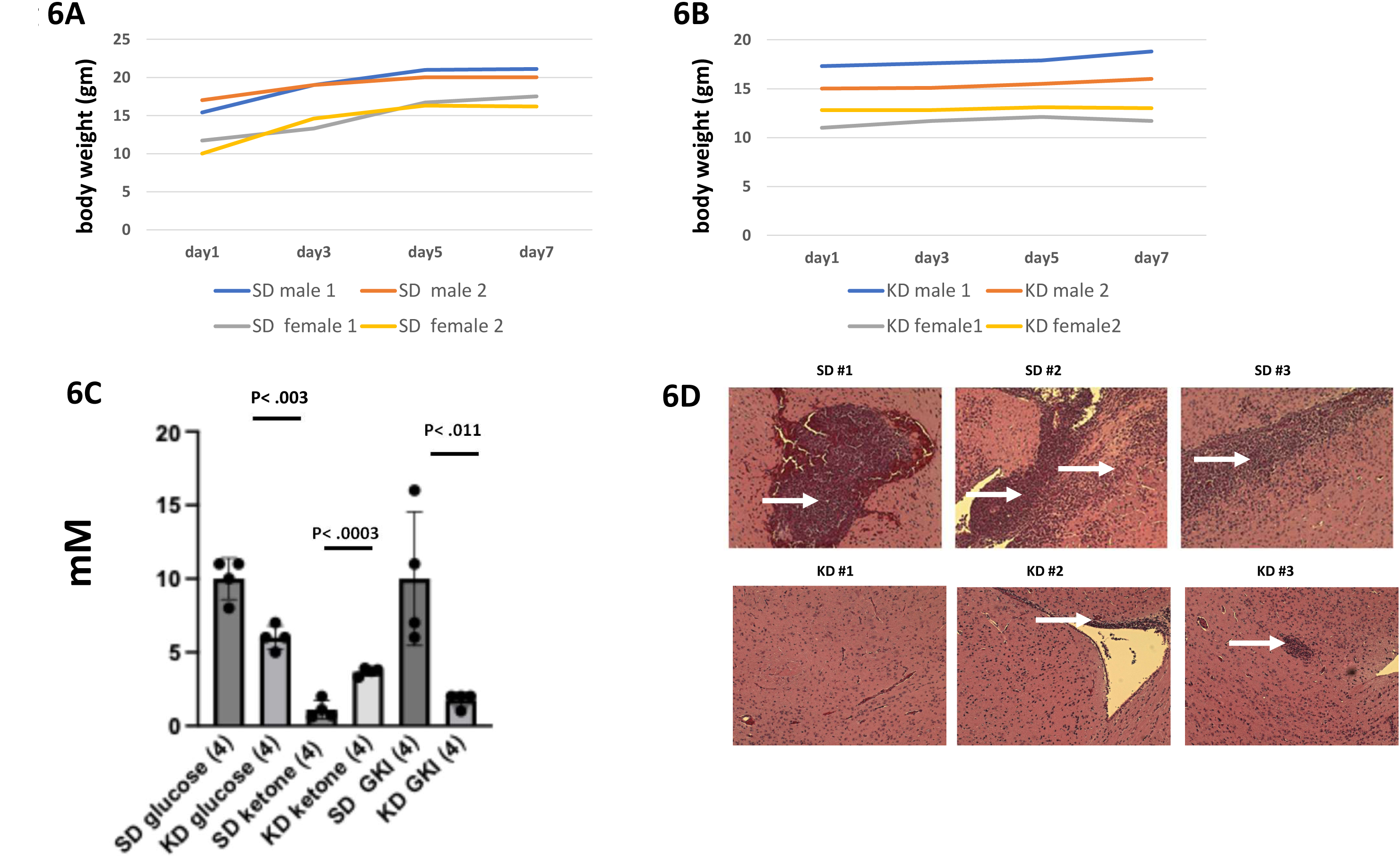
Influence of the KD on body weight, blood glucose, β-hydroxybutyrate, and the glucose ketone index (GKI) values in p21-25 mice on ketogenic diet and histology of the brains with VM-M3/luc cells. Juvenile mice (p25) were tested for the ketogenic diet tolerability and blood glucose and ketone level in the blood. **A.** Body weight increased in the mice receiving the SD but remained steady in the mice receiving the KD. **B.** A significant reduction of blood glucose is associated with a significant elevation of blood ketone levels in KD fed mice and accordingly. GKI. Values are expressed as the mean ± SEM and unpaired t-test determined the significance between the groups. **C**. Brain tissue histology (H&E staining) was evaluated 14 days after tumor implantation. The low power (10x) images on the top show many VM-M3/luc cells invading into normal appearing brain tissue in a standard diet fed mice (white arrows). Images on the bottom show a reduction of tumor cells with less invasion in KD-fed mice. 3 independent mice brains were fixed, processed and imaged.

Blood glucose levels were significantly lower in the mice fed the KD (5-7 mM) than in the mice fed the SD (8-11 mM). This significant reduction of blood glucose in the KD-fed mice was associated with a significant elevation of blood β-OHB levels, i.e., 3.3-3.9 mM in the KD-fed mice compared to 0.7-2.0 mM in the SD-fed mice. Accordingly, the GKI was significantly reduced in KD-fed mice compared to that of SD-fed mice (**Figure 6C**). These findings further indicate that KD therapy can reduce GKI values and tumor invasion compared to SD alone. Histological analysis revealed that VM-M3 glioma cell invasion was markedly less in the brains of the KD-fed mice than in the brains of SD-fed mice (**Figure 6D**). These findings suggest that ketogenic metabolic therapy could be therapeutic for children with HGG.

### MBZ and CPI-613 reduce the proliferation of human pediatric GBM cell, SF-188/luc *in vitro*

SF-188 is an important cellular model for pediatric human glioblastoma [69, 70]. To study the effect of MBZ and CPI-613 on the proliferation of human SF-188/luc, the cells were seeded and treated the same way as described above for VM-M3 cell (Figure 3A and 4A). DON was used as a positive control. SF-188/luc cells (1.0 × 10^5^) were seeded in 24-well plates in complete medium as described before. After 24 hours, the cells were rinsed with PBS and treated with MBZ (1.0 μM) and CPI-613 (100 μM) in basal media for 24 and 48 hours. Bioluminescence was significantly lower for both MBZ and CPI-613-treated SF-188 cells/luc after 24 hours (**Figure 7A**). Interestingly, both MBZ and CPI-613 treated cells showed an increase in caspase-3 expression, similar to what was observed in VM-M3 cells, suggesting involvement of apoptotic pathways (**Figure 7B-C**).

**Figure 7.**
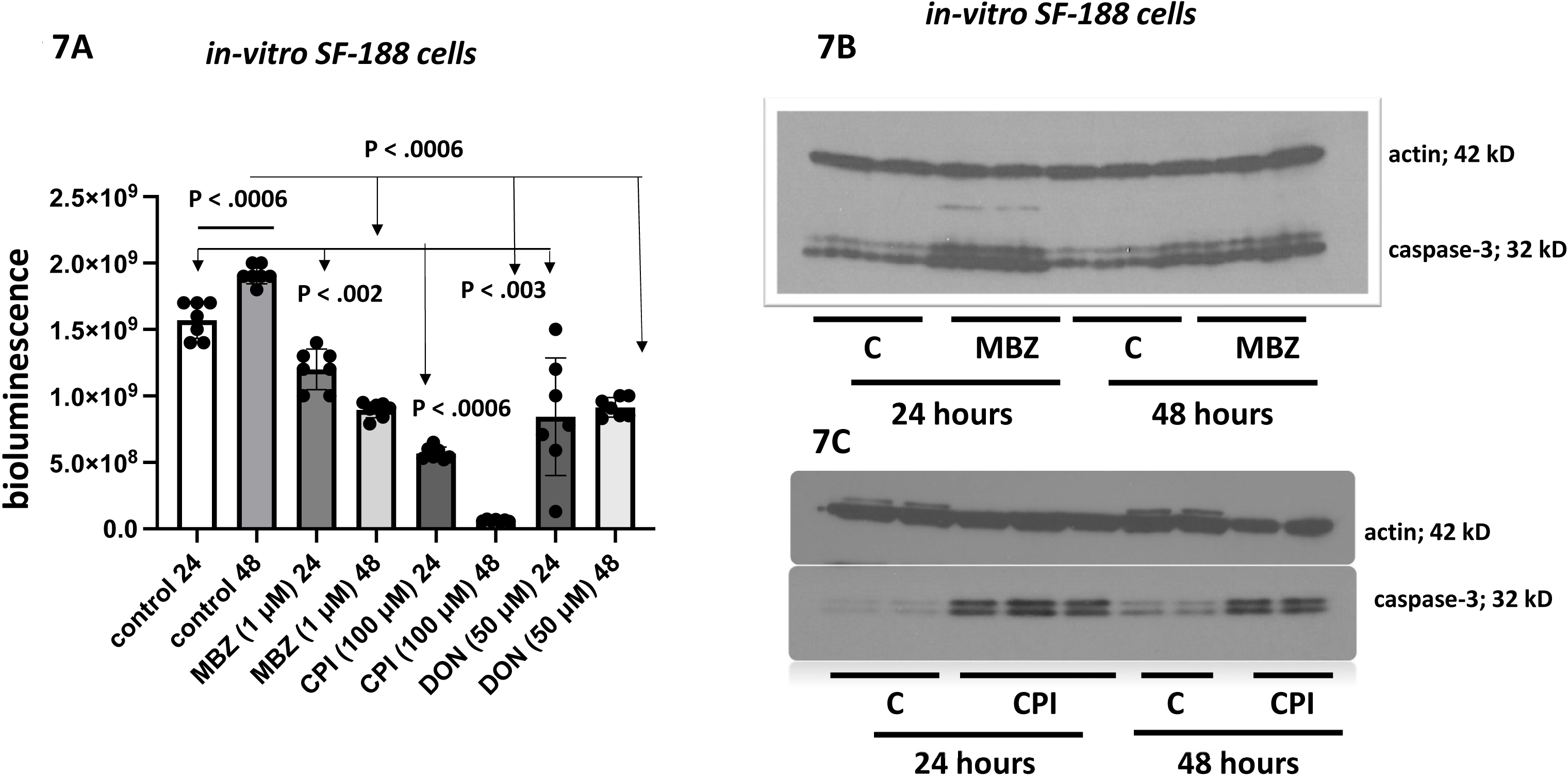
Effects of MBZ and CPI-613 (devimistat) on the progression of human pediatric SF-188 /luc cells *in vitro* SF-188 /luc cells were treated with MBZ (1μM) and CPI-613 (100 μM) for 24 and 48 hours while grown *in vitro*. **A**. Both drugs reduced the bioluminescence significantly in the SF-188 /luc cells in a time dependent manner (24 and 48 hours). The study was repeated, and the values are expressed as the mean ± SEM and Mann-Whitney test to determine the significance between the groups. **B and C.** MBZ (1μM) and CPI (100 μM) induced apoptosis in SF-188/luc cells indicated by caspase-3 activation in both 24 and 48 hours.

## Discussion

Our study is the first to show a robust therapeutic benefit of combining MBZ and CPI-613 with the KD for invasive HGG in a preclinical model. This approach highlights the value of diet–drug therapy in targeting glucose and glutamine metabolism. Previously, we found that KD plus DON prolonged survival in adult VM/Dk mice with VM-M3 gliomas, though without spinal cord invasion [33]. In contrast, juvenile (p21–25) mice with VM-M3 implants showed extensive spinal cord and meningeal invasion, consistent with the mesenchymal origin of these cells. Such invasion is absent with non-metastatic VM-NM1 or CT-2A gliomas of neural stem cell origin [18, 20, 61, 71]. These findings suggest that the juvenile CNS microenvironment is especially permissive for mesenchymal HGG invasion, making the VM-M3 model in p21 mice highly relevant to pediatric disease.

We also showed for the first time that the anti-parasitic drug MBZ could target the glutaminolysis pathway in VM-M3 glioma cells, which was linked to improving survival of the mice. Moreover, our findings showed that the PDH/α-KGDH inhibitor, CPI-613, was therapeutic against VM-M3 cell growth and invasion only when used together with the ketogenic diet. Ketogenic metabolic therapy, if administered appropriately in mice or humans, can be an effective press therapy for reducing inflammation, proliferation, and angiogenesis while at the same time enhancing the death of malignant glioma cells [28, 60]. Consequently, we administered MBZ and CPI-613 to the juvenile mice with the KD to improve bioavailability and reduce toxicity.

The p21 mice used in our study were administered the KD *ad-libitum* to help maintain body weight, which can be challenging in juvenile mice [72]. All the juvenile mice on KD achieved therapeutic ketosis within 3-4 days of diet treatment. The brain histology data showed that tumor growth was significantly less in the KD-fed mice than in the SD-fed mice. From our experience, the anti-cancer therapeutic effect of the KD is best when used against naturally arising tumors grown in their syngeneic host using fat: protein + carbohydrate ratios of 3:1 or 4:1. These ratios can also reduce the glucose ketone index (GKI) to 2.0 or below with modest to minimal body weight loss. Studies not using natural brain tumor models or adhering to these diet criteria often do not show anti-cancer effects of ketogenic metabolic therapy (KMT) [73–76]. The therapeutic success of KMT comes largely from optimizing the macronutrient and micronutrient composition of the KD to achieve the therapeutic GKI values [19, 60, 68]. As the basal metabolic rate is slower in humans than in mice, the therapeutic effects of the KD may be potentially more powerful in children with HGG than in mice with HGG. Glutamine-driven mesenchymal cells are the most invasive malignant cells in HGG [27, 33]. The KD will be less able to restrict growth of these glutamine-dependent neoplastic cells than those cells more dependent on glucose. In this situation, the pulsing of glutamine targeting drugs can be used safely with a nutritionally balanced GKI controlled ketogenic diet (the press) to maximize the therapeutic effect of drug interventions [60].

Previous studies showed that the bioavailability of MBZ was improved when administered with high fat meals [42, 44]. Additional strategies used to increase drug bioavailability in tumors include alternative formulations of MBZ with vegetable oils, altering the crystalline structure, and PEGylation [77–80]. Oral administration of MBZ (50 mg/kg mixed with sesame oil) daily in syngeneic and orthotopic glioma models in adult mice significantly extended mean survival up to 63% [35]. We showed that the KD used alone improved the survival of the tumor-bearing juvenile mice up to 100% over the SD-fed mice. Further improvement in survival was seen when we administered MBZ pulsing (3x a week) together with the KD. The cocktail of MBZ and DON with ketogenic diet extended the survival over 3-fold. We previously showed that DON is apowerful glutamine targeting drug that significantly inhibits VM-M3 tumor progression [33]. Consequently, we included a DON-treated group as a positive control in our current study. There are currently DON prodrugs in clinical trials for different cancers, but none to our knowledge are in use with KD following a press-pulse implementation [81].

We previously showed that the KD could facilitate delivery of structurally different small molecules through the blood brain barrier [33, 67]. DON delivery to the orthotopic VM-M3 tumor was three-fold greater when administered with a calorie restricted KD mice than when administered with high-carbohydrate mouse standard diet [33]. Moreover, MBZ must be administered with a fat-containing diet, either mixed into the food or delivered by gavage. Consequently, we did not include a SD + MBZ mouse group in this study. The clear presence of MBZ in the brains of mice treated with KD + MBZ further validates effective delivery. Administration of MBZ and DON with the KD allowed for lower dosing or pulsing, thus mitigating issues of drug toxicity. Better overall mouse survival without observable toxicity was achieved when both MBZ and DON were administered with the KD. This information would be especially important when translating this therapeutic strategy to the clinic, as avoidance of any type of toxicity would be of upmost importance to children with HGG.

The therapeutic action of MBZ is due in part to inhibition of tubulin polymerization [35, 82]. Tubulin polymerization is an ATP driven process in parasites [83]. Previous reports have suggested that inhibition of carbohydrate metabolism might also account in part for the anti-parasitic action of MBZ [84, 85]. We now show that MBZ could block the glutamine utilization in VM-M3 cells *in vitro* and could inhibit the expression of glutaminase C in *in vivo* tumor tissue. Glutaminase C expression is directly related to the malignancy of different tumor cells and thus glutamine metabolism is elevated in many cancers [65, 86, 87]. Metabolic cooperation of both glucose and glutamine is essential for the life cycle of both parasites and tumor cells [88]. We described how glucose and glutamine are two major fuels that drive the cytoplasmic and mitochondrial substrate level phosphorylation necessary for tumor cell survival and growth [31, 63, 89]. Further study on the human pediatric glioma cell line SF-188 showed that MBZ is equally effective in reducing cell proliferation *in vitro* as in the mouse cell lines VM-M3 and CT-2A. We are not aware of any ideal pediatric GBM xenograft model which shows robust invasion and growth like VM-M3 cells, especially to test metabolic therapy *in vivo*. Moreover, in preclinical studies, the natural host plays a crucial role in evaluating therapeutic efficacy. PDX models may present challenges for tumor implantation into immunocompetent, juvenile mouse brain. Therefore, further studies will be needed to test similar diet-drug approaches in xenograft or genetically engineered humanized models, with therapeutic effect clearly demonstrated in a syngeneic context.

Our study involved the evaluation of CPI-613 (devimistat) as a potential therapy for pediatric HGG. CPI-613 targets the pyruvate dehydrogenase (PDH) and α-ketoglutarate dehydrogenase enzymes necessary for tumor growth and is in clinical trials for pancreatic cancer in combination with other drugs [49, 54]. A previous study showed that CPI-613 could reduce TCA cycle metabolites in an adult mouse brain tumor model [55]. We found a significant dose-dependent inhibitory effect of CPI-613 on VM-M3 cells *in vitro.* These findings justified an evaluation of CPI-613 on the *in vivo* growth of the VM-M3 glioma in p21 mice. Remarkably, we found that CPI-613 was effective in reducing VM-M3 tumor growth and invasion only when administered with the ketogenic diet. CPI-613 had no significant effect on VM-M3 tumor growth when administered alone in the standard (SD) mouse chow diet. In addition to improving the survival of p21 VM/Dk mice bearing the VM-M3 glioma, the combination of KD and CPI-613 also significantly improved overall survival in p21 C57BL/6 mice bearing the CT-2A neural stem cell tumor. Our findings are also consistent with previous findings showing that a KD is therapeutic against human glioma stem cell tumors [90]. The mechanism by which KD enhances delivery of small molecule drugs through the blood brain barrier remains unclear. It can be challenging to isolate and purify water-insoluble drugs for LC-MS analysis from brain tissue. Further studies will be needed to explore this issue. OGDH, a target of CPI-613, shows significantly lower expression in the brain tissue of the KD + CPI group compared to the SD + CPI group, which further suggests that drug delivery may be more effective in the latter group. Additional quantitative studies are necessary to confirm this observation.

The ketogenic diet, MBZ, and CPI-613 are currently in clinical trials as single-agent interventions for different cancers in adults and children. The ketogenic diet is already well-recognized as an effective therapy for managing epilepsy in children. Importantly, Nebeling *et al*. first reported the therapeutic effect of a ketogenic diet on HGG in two pediatric patients and linked the glucose/ketone metabolism to their improved quality of life and overall survival [91]. Unfortunately, the study remained largely unrecognized for years despite evidence showing that the outcomes were significantly better than that from the standard of care. MBZ is in clinical trials for pediatric recurrent/non-responsive brain tumors to evaluate the safety and tolerability of different dosing regimens. Our preclinical results in the juvenile p21 mouse model could have a direct translational benefit to childhood brain cancer, if administered appropriately with the KD. We also expect that this therapy could be effective for managing childhood HGG when used either alone or in combination with non-toxic aspects of standard of care. the Additional preclinical studies are needed for CPI-613 in relevant brain tumor models before considering use in clinical trials for pediatric glioma. This study primarily focused on using metabolic therapy alone for childhood HGG to bypass the toxicity associated with standard care [25, 92]. However, this does not rule out the possibility that outcomes could be improved if used as an adjuvant treatment, especially for adult GBM. Further investigations are underway in our laboratory to improve metabolic therapy using combinatory regimens of MBZ, CPI-613, DON prodrug, and other anti-glutamine drugs with ketogenic diet for management of pediatric HGG. We believe our study will have a potential bench to bedside translation, especially for children with HGG.

### Conclusion/Summary/Significance/Limitation

This study demonstrates the potential of combining nutritional ketosis with targeted drug therapies to improve outcomes in childhood brain cancer. Implementing a ketogenic diet (KD) enhanced drug efficacy by reducing tumor invasion, slowing tumor growth, and extending survival in juvenile mice with high-grade gliomas. Notably, KD also permitted the use of lower drug dosages, thereby minimizing toxicity and reducing the risk of long-term side effects. The press-pulse strategy (simultaneously targeting glycolysis and glutaminolysis pathways) emerges as a promising, less toxic approach for managing aggressive pediatric brain tumors [60, 93]. These findings underscore the importance of integrating dietary interventions with pharmacological treatments and highlight the translational potential from bench to bedside, offering hope for improved survival and quality of life in children with HGG. The main limitation of this study is that the precise mechanism by which the ketogenic diet enhances drug delivery to tumor tissue remains unclear. Further *in vivo* analyses under different conditions will be required to address it. Ongoing studies are extending this approach to additional tumor models.

## Materials and Methods

### Mice

Juvenile mice (*p21-p25*) of the VM/Dk (VM) and C57BL/6J (B6) strain were used for this study. Studies were initiated one day after weaning. Males and females within and between litters were matched for age, sex and body weight. All mice used in this study were housed and bred in the Boston College Animal Care Facility using husbandry conditions as previously described [94]. All animal procedures and protocols were in strict accordance with the NIH Guide for the Care and Use of Laboratory Animals and were approved by the Institutional Animal Care Committee at Boston College under assurance number A3905-01.

### Syngeneic HGG glioma model grown in juvenile mice

The (p21-p25) mice were anaesthetized with isoflurane (5% in oxygen). The tops of the heads were shaved and disinfected with ethanol and a small incision was made in the scalp over the midline. A small indentation was made in the skull over the right parietal region behind the coronal suture and lateral to the sagittal suture using an 18G needle. VM-M3 tumor cells (approx. 1.0 × 10^5^ in 5 µL PBS) were implanted in the right cerebral cortex approximately 1.5–2.0 mm deep using a Hamilton syringe. Betadine, a topical antiseptic, was applied before the skin flaps were closed with 7 mm reflex clips. The mice were placed in a warm room (24°C) until they were fully recovered. The procedure confirms 100% recovery within a few hours of implantation. CT-2A tumor cell (approx. 1.0 × 10^5^ in 5 µL PBS) were implanted in juvenile C57/BL6 mice (p21-p25), respectively, using the same procedure described above for VM-M3 cells.

### Tumor cell lines

The VM-M3 invasive glioma used in this study originally arose spontaneously in the cerebrum of an adult male mouse of the VM/Dk inbred strain. Spontaneous brain tumors arise more frequently in this strain than in other mouse strains [95]. A cell line was prepared from the tumor as described previously [20]. The VM-M3 tumor manifests all the invasive characteristics seen in human GBM [61]. The CT-2A tumor was originally produced from implantation of 20-methylcholanthrene into the cerebral cortex of a C57BL/6J mouse and was broadly classified as a poorly differentiated highly malignant anaplastic astrocytoma or neural stem cell tumor and a cell line was produced from this tumor as described [71, 96]. Like the VM-M3 tumor, the VM-NM1 tumor arose spontaneously in the cerebrum of an adult mouse of the VM/Dk inbred strain [20]. In contrast to the highly invasive mesenchymal VM-M3 tumor cells, the VM-NM1 glioma cells are non-invasive and express multiple markers of neural stem cells like that seen in the CT-2A cells [20]. SF-188 cells were obtained from Sigma (Cat. No. SCC282). All tumor cell lines were transfected with a lentivirus vector containing the firefly luciferase gene under control of the cytomegalovirus promoter (VM-M3/Fluc, gift from Miguel Sena-Esteves, UMass Medical School). This transfection allowed the cells to be tracked in the brain using bioluminescent imaging. All tumor cells were cultured in DMEM containing 25.0 mM glucose and 4.0 mM glutamine (Sigma D-5796) supplemented with 10% fetal bovine serum (FBS, Atlas Biologicals, CO). Basal media is also DMEM (Sigma D-5030) but without supplemented glucose and glutamine.

VM-M3 cells (1.0 × 10^5^ cells) were seeded in 24-well plates using DMEM and 10% FBS. After 24 hours, the cells were rinsed with PBS and then grown in serum free basal media with 12.0 mM glucose and 2.0 mM glutamine. The cells were cultured for another 24 hrs in MBZ (vehicle DMSO) or DON using different drug concentrations. DON was used as a positive control for these experiments as our previous studies showed that DON targets the glutaminolysis pathway by inhibiting multiple glutaminases [97, 98]. Cells were collected at 6 and 12 hours for LC-MS analysis of metabolites and bioluminescence.

### Dietary/Drug Regimens and Body Weight

All juvenile mice received the standard diet (SD) for a day or two after weaning and prior to initiation of the study. Tumor cells were implanted on day zero. Upon implantation of the tumor, mice remained on the SD for 6 days until an *in vivo* image was performed to confirm the tumor take for all mice. Mice were assigned to diet and drug groups on day 6. Male and female juvenile mice were both used for each group. Mice receiving the standard diet (*LabDiet* 5P00-Prolab^®^ RMH 3000) *ad libitum* for the duration of the study. Ketogenic diet (KetoGEN, Medica Nutrition, Canada) were given *ad-libitum* so that the juvenile mice can maintain the body weight for this short-term study. The percent nutritional breakdown of KetoGEN and Standard diet as described (Supplement Table 1). MBZ (100 mg/kg body weight, Sigma-Aldrich, St. Louis, MO) was mixed thoroughly in the KD and administered to the mice for 3 consecutive days/wk. Food intake was monitored daily and given fresh to approximate the amount of drug consumed for each mouse. The KD, without MBZ, was administered alone on the off days. For those mice that received DON injections, a fresh stock of DON was prepared and diluted to an appropriate concentration in PBS and was administered intraperitoneally (i.p.). The DON stock solution in PBS was stored at -20°C for the duration of the study and mice received 100-150 μL injections of 1.0 mg/kg. DON was injected one or two times a week depending on the overall health and tumor progression of the mice. Some doses were skipped if the mice appeared lethargic or if body weight loss exceeded 1.0 g from the previous day. Studies were terminated at the time of morbidity for the control groups. CPI-613 (AdooQ Biosciences, Irvine, CA) was injected i.p. 0.5-1.0 mg/kg body weight once or twice in a week. CPI-613 stock was made in DMSO solution and fresh 100x diluted in PBS before injection in mice. Similar dilution of DMSO was injected in the control group. We did not administer the drugs using gavage in the juvenile mice as gavage-induced adverse effects and hyperglycemia would have confounded data interpretation [99, 100].

### Bioluminescence Imaging

The AMI HT imaging machine (Spectral Instruments Imaging, Tuscon, AZ) was used to record the bioluminescent signal from the labeled tumors as we previously described. Briefly, for *in vivo* imaging, mice received an i.p. injection of D-lucifierin (50 mg/kg) in PBS. Imaging times ranged from 1 to 5 min, depending on the time point. For *ex vivo* imaging, brains were removed and imaged in 0.3 mg D-luciferin in PBS. The IVIS Lumina cooled CCD camera system was used for light acquisition. Data acquisition and analysis was performed with Living Image software (Caliper LS).

### Survival study in juvenile mice

Juvenile mice were implanted and received the diet/drug regimen as described above. Mice were fed *ad libitum*, and body weight was monitored daily. Diet and drug were administered as described above. Any mouse from any group that lost more than 20% of its body weight or showed signs of illness was considered censored.

### Histology and Immunohistochemistry

Brain tumor, spinal cord, and liver tissues were fixed in 10% neutral buffered formalin (Sigma-Aldrich) and embedded in paraffin. Tissues were sectioned at 5 µm and were stained with hematoxylin and eosin (H&E) performed at the Harvard University Rodent Histopathology Core Facility (Boston, MA). Tissue sections were examined by light microscopy using either a Zeiss Axioplan 2 or Nikon SMZ1500 light microscope. Images were acquired using SPOT Imaging Solutions (Diagnostic Instruments, Inc., Sterling Heights, MI) cameras and software. All histological sections were evaluated at the Harvard University Rodent Histopathology Core Facility. For Ki67 staining, immunohistochemistry was performed using the same procedure as we previously described (31).

### Liquid Chromatography Mass Spectrometry Analysis of Metabolomics

VM-M3/luc cells were seeded (1.0 × 10^5^) in 6-well plates. After 24 hours the cells were treated with MBZ (1.0 µM) for 2, 6, or 12 hours in basal media with no FBS (12 mM glucose and 2.0 mM glutamine). After removing the media, 1.0 mL of ice cold methanol:water (80:1) mixture was added to each well and immediately transferred to -80°C for 15 minutes. Plates were taken out at different time points and placed on ice, scraped, and the extract transferred to Eppendorf tubes. The tubes were vortexed and centrifuged at 10,000 x g for 10 minutes. The supernatant was transferred to a new tube and centrifuged at 30°C. Cell extract was reconstituted in 100mL of methanol and then filtered through nylon before injecting into the LC-MS instrument (Agilent 6220 TOF, BPGbio, Framingham, MA).

### Tumor and Tissue Analysis Using MALDI Mass Spectrometry and Microscopy

#### Preparation of tissue for MALDI MSI and microscopy

Mouse brains were dissected and snap frozen in liquid nitrogen and stored at -80°C. Coronal cryosections (10 μm thickness) were obtained using a cryostat (CryoStar NX50, Thermo Scientific, Kalamazoo, MI) and thaw-mounted on indium tin oxide (ITO) slides. Additional serial sections were used for MALDI MSI and hematoxylin and eosin (H&E) staining. Optical microscopy scan images were collected at 5 μm pixel size using a slide scanner (LI-COR Odyssey M, Lincoln, NE).

#### Preparation and application of MALDI matrix

A 2,5-dihydroxybenzoic acid matrix (80 mg/mL in 70:30 methanol: water with 0.1% TFA and 1% DMSO) was used to detect mebendazole. The matrix was applied using a TM Sprayer (HTX M3 Technologies, Chapel Hill, NC) run at 0.18 mL/min flow rate, at total drops of 1200 mm/min, and using 10 psi nitrogen pressure, 75 °C nozzle temperature and 3 mm track spacing, at 10 passes.

MALDI MRM MS imaging: Sections were imaged on a timsTOF fleX mass spectrometer (Bruker Daltonics, Billerica, MA) in positive ion mode using multiple reaction monitoring (MRM) over the m/z range from 50–1600. A mebendazole standard was infused via ESI, allowing optimization of the MRM parameters: ion funnels, quadrupole, collision energy, and TOF focus. The optimal transition was 296.103 → 264.077 (3 m/z isolation width, 15 eV collision energy) related to [C□□H□□N□O□+ H] and [C□□H□N□O□+ H]□. The method was adapted for MALDI and tuned using an Agilent tune mix (Agilent Technologies, Santa Clara, CA). Imaging was performed at 50 μm pixel size and 10,000 Hz laser repetition rate and 2,000 shots per pixel.

### Visualization of data

MSI data were analyzed by SCiLS Lab software (version 2025b Premium, Bruker Daltonics) with total ion current (TIC) normalization.

#### Ultra-high performance liquid chromatography mass spectrometry (UHPLC-MS) analysis of CPI-613 in serum

Mice were injected with 1.0 mg/kg CPI-613 at 30-45 min before collection of the blood.The blood samples were collected, sera were separated and stored at -80 °C. CPI-613 wasextracted from 50µL aliquots following the addition of 300µL isopropanol, vortexing for1min, and centrifuging at 14,000g for 10min. The supernatants were transferred intoLCMS vials and stored at 4°C until analysis. UHPLC-MS was performed using anAgilent 1290 LC equipped with a Waters Acquity Premier CSH column (15cm x 2.1mm x1.7µM) kept at 55°C, coupled to a Thermo Q-Exactive Plus mass spectrometer. Thebinary multi-step gradient was composed of A: 60/40 acetonitrile/water and B: 90/10isopropanol/acetonitrile, both with 0.1% formic acid, 10mM ammonium formate and0.1µM reserpine (lock mass) with a flow of 230µL/min. The mass spectrometer wasoperated in full-scan mode, 17,500 resolution, scan range m/z 120 – 1600, with fastpolarity switching and spray voltages +3.5kV (positive) and -2.5kV (negative). Spectrawere automatically centroided and lock-mass corrected during acquisition.

#### Oxygen Consumption Assay

Oxygen consumption was measured continuously using the Resipher instrument (Lucid Scientific, Atlanta, GA) in 96-well Falcon flat-bottom plates as described previously [32, 101]. Cells were seeded at a density of 2.0 × 105 cells/well. The sensing lid and Resipher device were placed on top of the 96-well plate for five minutes prior to the start of the experiment. Experimental media containing MBZ and CPI-613 was added in different set of experiments. The sensing lid was placed on top of the Falcon plate and put into the HEPA 100 incubator (Thermo Fisher Scientific, Waltham, MA) maintained at 5% CO2 and 88% relative humidity. Oxygen consumption rate (OCR) was monitored in the incubator for 6 hours. Data was analyzed using the Resipher web application.

#### Blood glucose and ketone measurements

All mice were fasted for 2 hours to stabilize blood glucose levels before blood collection. Blood glucose and ketone levels were measured using the Keto-Mojo monitoring system (Keto-Mojo, Napa, California). Whole blood from the tail was placed onto the glucose or ketone strip. The keto-mojo meter was used to determine the mmol levels of glucose and β-hydroxybutyrate in the blood. The Glucose Ketone Index (GKI) was determined as we previously described [19, 68].

#### Serum Alanine Transaminase assay

Serum ALT activity was measured using a colorimetric assay kit (Cayman Chemical, MI, USA). Briefly, serum was separated from blood samples and stored at −80 °C until analysis. The assay was performed in a 96-well plate using 20 µL of serum per well, with each sample run in triplicate. The manufacturer’s protocol was followed, and kinetic absorbance was measured using a SpectraMax M5 plate reader (Molecular Devices, USA).

#### Western blot analysis of glutaminase, OGDH, and caspase-3 protein expression

Right side of the frozen brain tissues where tumor cells were implanted were used for protein expression. Tissues were homogenized in ice-cold lysis buffer containing 20 mmol/L Tris-HCl (pH 7.5), 150 mmol/L NaCl,1 mmol/L Na_2_EDTA, 1 mmol/L EGTA, 1% Triton, 2.5 mmol/L NaPPi,1 mmol/L α-glycerophosphate, 1 mmol/L Na_3_PO_4_, 1 µg/mL leupeptin, and 1 mmol/L phenylmethylsufonyl fluoride. Lysates were transferred to 1.7 mL Eppendorf tubes, mixed on a rocker for 1 h at 4°C, and then centrifuged at 8,100 x g for 20 min. Supernatants were collected and protein concentrations were estimated using the Bio-Rad detergent-compatible protein assay. Approximately 100 µg of total protein from each tissue sample was denatured with SDS-PAGE sample buffer {63 mmol/L Tris-HCl (pH 6.8), 10% glycerol, 2% SDS, 0.0025% bromophenol blue, and 5% 2-mercaptoethanol} and was resolved by SDS-PAGE on 4% to 12% Bis-Tris gels (Invitrogen). Proteins were transferred to a polyvinylidene difluoride immobilon TM-P membrane (Millipore) overnight at 4°C and blocked in either 5% nonfat powdered milk or 5% bovine serum albumin in TBS with Tween 20 (pH 7.6) for 1 to 3 h at room temperature. Membranes were probed with primary antibodies (glutaminase C, OGDH, Abcam, UK; Caspase-3, Cell Signaling, MA) overnight at 4°C with gentle shaking. The blots were then incubated with the appropriate secondary antibody (anti-rabbit) for 1 h at room temperature and bands were visualized with enhanced chemiluminescence. Each membrane was stripped and re-probed for β-actin (Cell Signaling, MA) as an internal loading control.

#### Statistics

Tumor bioluminescence *in vitro*, *in vivo*, and *ex vivo* data were analyzed using the one-way analysis of variance (ANOVA) followed by Mann-Whitney test or by a student’s t test. In each figure, error bars are mean ± SEM and *n* is the number of individual mice analyzed. The Survival studies were plotted on a Kaplan-Meier curves were determined using Prism (GraphPad Software, Inc., San Diego, CA) and significance was analyzed using the log-rank test.

## Abbreviations

DMSO: dimethyl sulfoxide
PBS: phosphate buffered saline
PDH: pyruvate dehydrogenase
α-KGDH: α-ketoglutarate dehydrogenase
OGDH: 2-oxoglutarate dehydrogenase
OxPhos: oxidative phosphorylation
Na_2_EDTA: Disodium ethylenediaminetetraacetate dihydrate
Na_3_PO4: trisodium phosphate
FBS: fetal bovine serum
MBZ: mebendazole
DON: 6-diazo-5-oxo-L-norleucine
KD: ketogenic diet
SD: standard diet
KMT: ketogenic metabolic therapy
ATP: adenosine triphosphate
TCA: tricarboxylic acid;
SEM: standard error of mean

## Declarations

### Ethics approval and consent to participate

N/A

### Consent for publication

All authors on this manuscript have provided consent for publication.

### Availability of data and material

The authors declare that all data supporting the findings of this study are available upon request from the corresponding author. The source data for all figures are provided as a source data file.

### Competing interests

The authors declare no competing interests.

## Acknowledgements

We thank Children with Cancer UK (Grant Ref 19-313), Foundation for Metabolic Cancer Therapies, Dr. Edward Miller, Dr. Joseph Maroon, the Corkin Family Foundation, Kenneth Rainin Foundation, the Delaware County Special Deputies Benevolent Fund, the Broken Science Initiative, and the Boston College Research Expense Fund for their support. We thank Bret Judson, director of core imaging facility, BC for his expertise and help in capturing images for publication.

## Authors’ contributions

P.M. conceived, designed, and performed the studies, analyzed the data, and wrote the manuscript. T.N.S. reviewed and edited the manuscript, acquired funding, and supervised the project. S.S., S.K., B.G., J.H. and M.A.K. performed LCMS experiments and analysis. A.C. and J.M. helped in animal experiments. N.T., D.C.L., and T.D. helped in data analysis and editing of the manuscript and figures. R.T.B. helped in animal histopathology analysis.

---

**Extended figure 2a.**
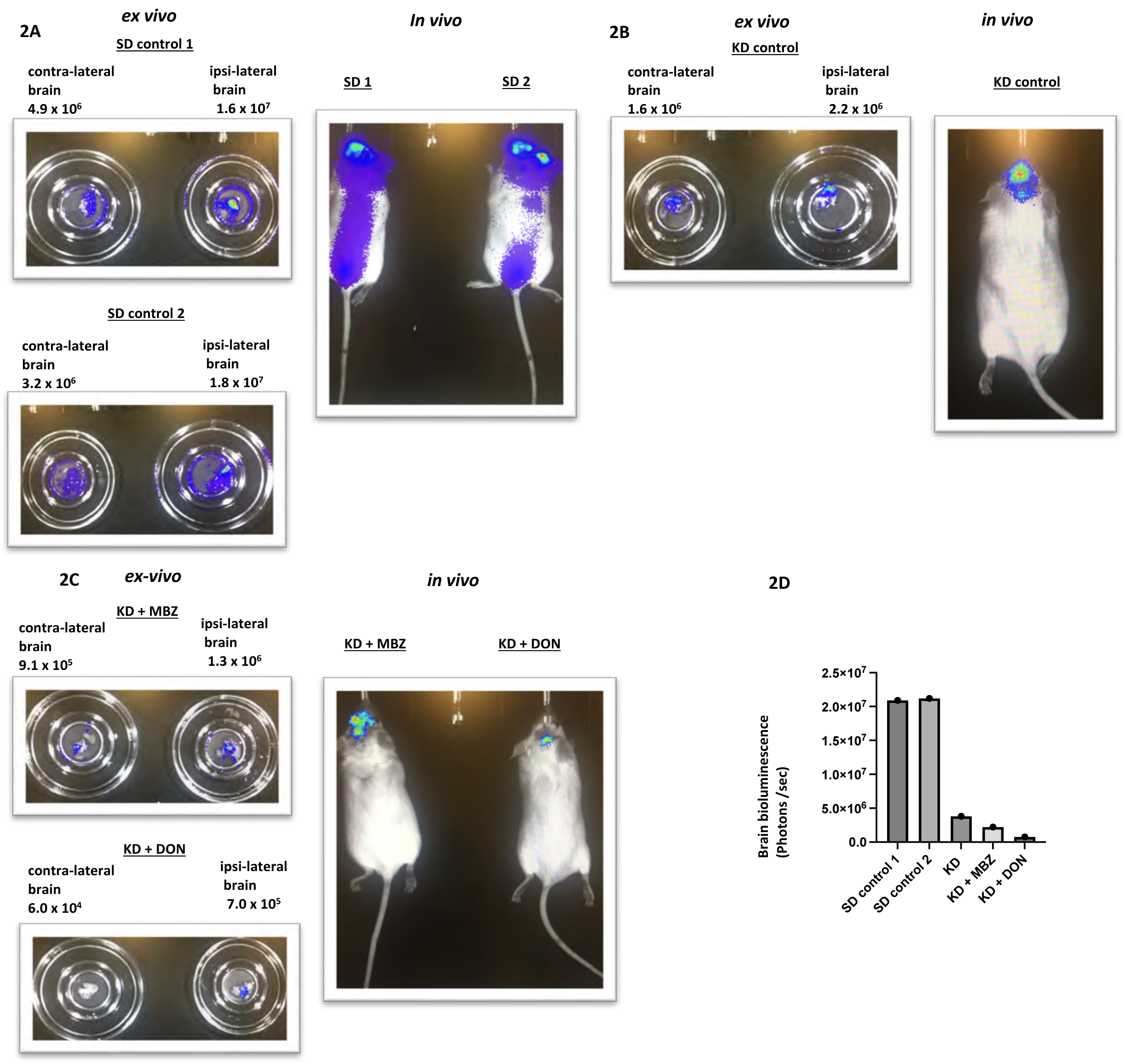
Effects of MBZ used with KD on orthotopically implanted VM-M3/luc glioblastoma cells in the brain of p20-25 VM/Dk mice. A preliminary study **A.** Analysis of mice receiving the control standard mouse chow diet as described in results. The “right” ipsilateral side of the brain (tumor implanted side) showed significant bioluminescence. Bioluminescence detected on the “left” contralateral side is indicative of significant distal hemispheric invasion of the tumor cells (*ex vivo* analysis). **B.** Analysis of mice receiving the Ketogenic diet (KD). No apparent spread of VM-M3/luc to the spinal column was detected on *in vivo* analysis. Bioluminescence in the ipsilateral and contralateral brain was noticeably lower in the KD-fed mice than in the SD control mice. **C.** Analysis of mice receiving either KD + MBZ or KD + DON. Brain bioluminescence was noticeably lower in these two groups than in either the SD control or the KD alone groups. Remarkably, no bioluminescent tumor cells were detected on the “left” contralateral brain side of the mouse treated in the KD + DON mice indicating that this treatment significantly reduced growth and distal invasion of the VM-M3/luc tumor cells. **D**. Bioluminescence *ex vivo* values from 5 mice mentioned above.

**Extended figure 2c-d.**
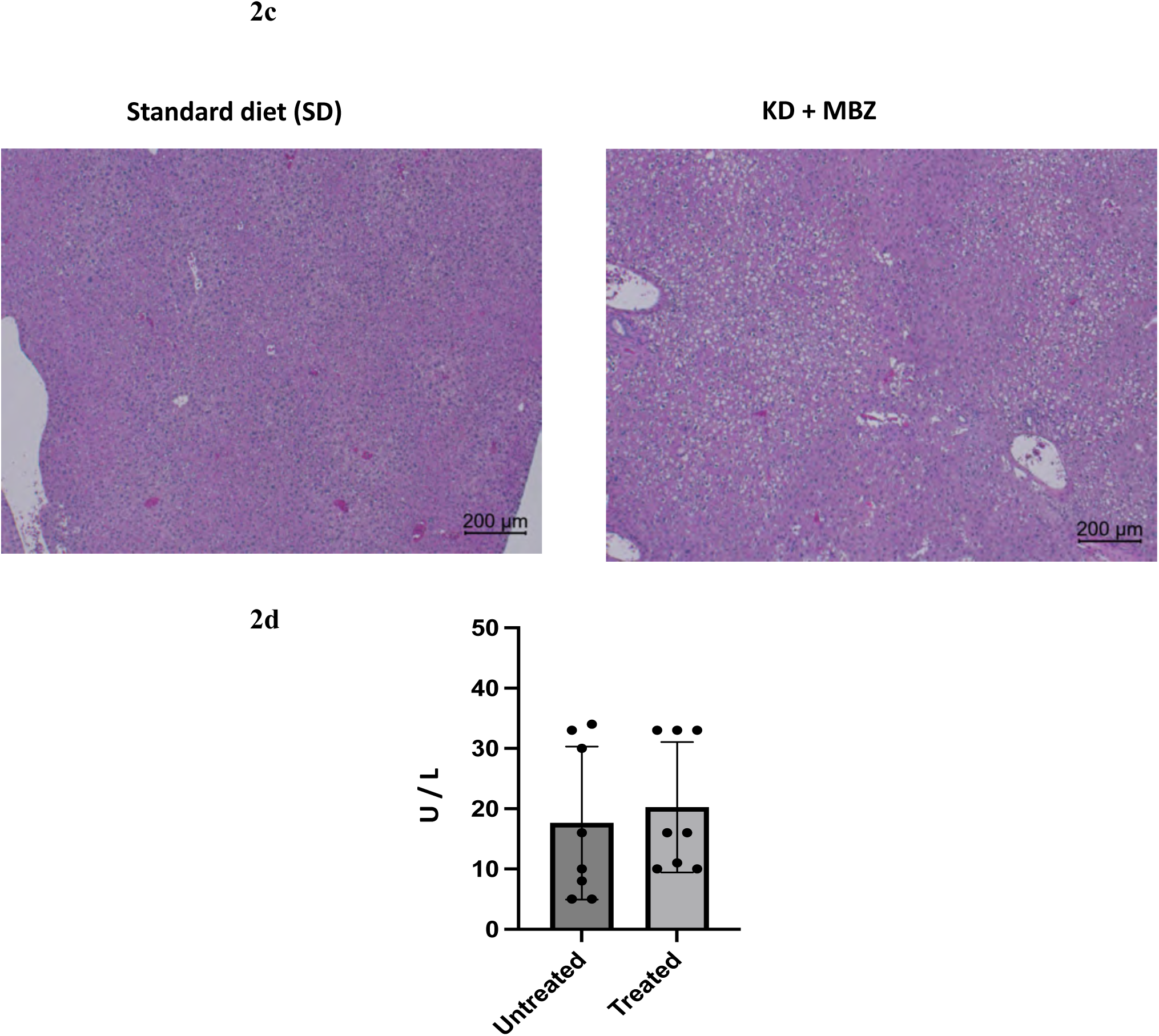
Effects of MBZ used with KD on orthotopically implanted VM-M3/luc glioblastoma cells in the brain of p20-25 VM/Dk mice. The p20 mice were implanted with VM-M3/luc tumor cells into the cerebral cortex and implemented diet and MBZ as described in results. At termination, liver and blood samples were collected for toxicity assessment. **c.** Liver histology (H&E) from the KD + MBZ group shows no evidence of drug-related toxicity (e.g., enlarged hepatocytes, necrosis, or immune cell infiltration) (Right) compared with liver from the untreated SD group (Left). Mild fat deposition was observed in the KD group, consistent with ketogenic diet effects. **d.** Serum alanine transaminase (ALT) activity levels (unit per liter) were comparable between groups, further indicating lack of hepatic toxicity.

**Extended figure 3a.**
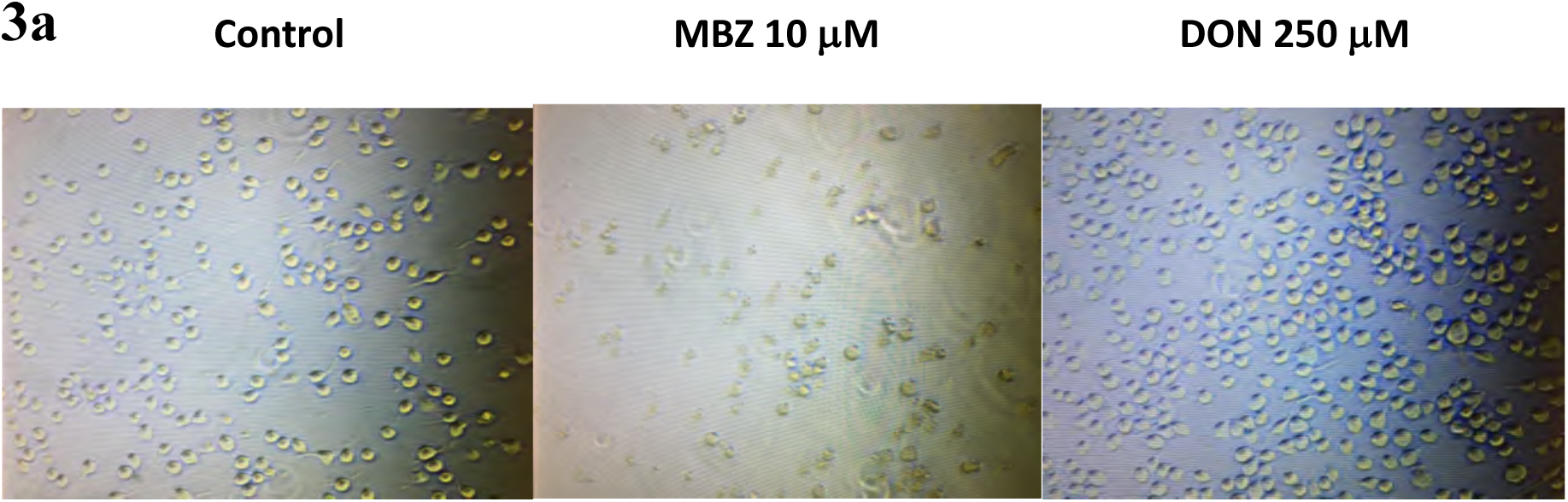
MBZ targets both glutaminolysis and glycolysis pathways in cultured VM-M3/luc cells. VM-M3/luc cells treated with high dose of MBZ and DON while cultured *in vitro* for 24 hours, following the described protocol. DON was included as a positive control. Brightfield images demonstrate that MBZ effectively killed VM-M3/luc cells at the highest dose. In contrast, DON reduced the proliferation rate of VM-M3/luc cells but did not induce cell death at the highest dose.

**Extended figure 3b.**
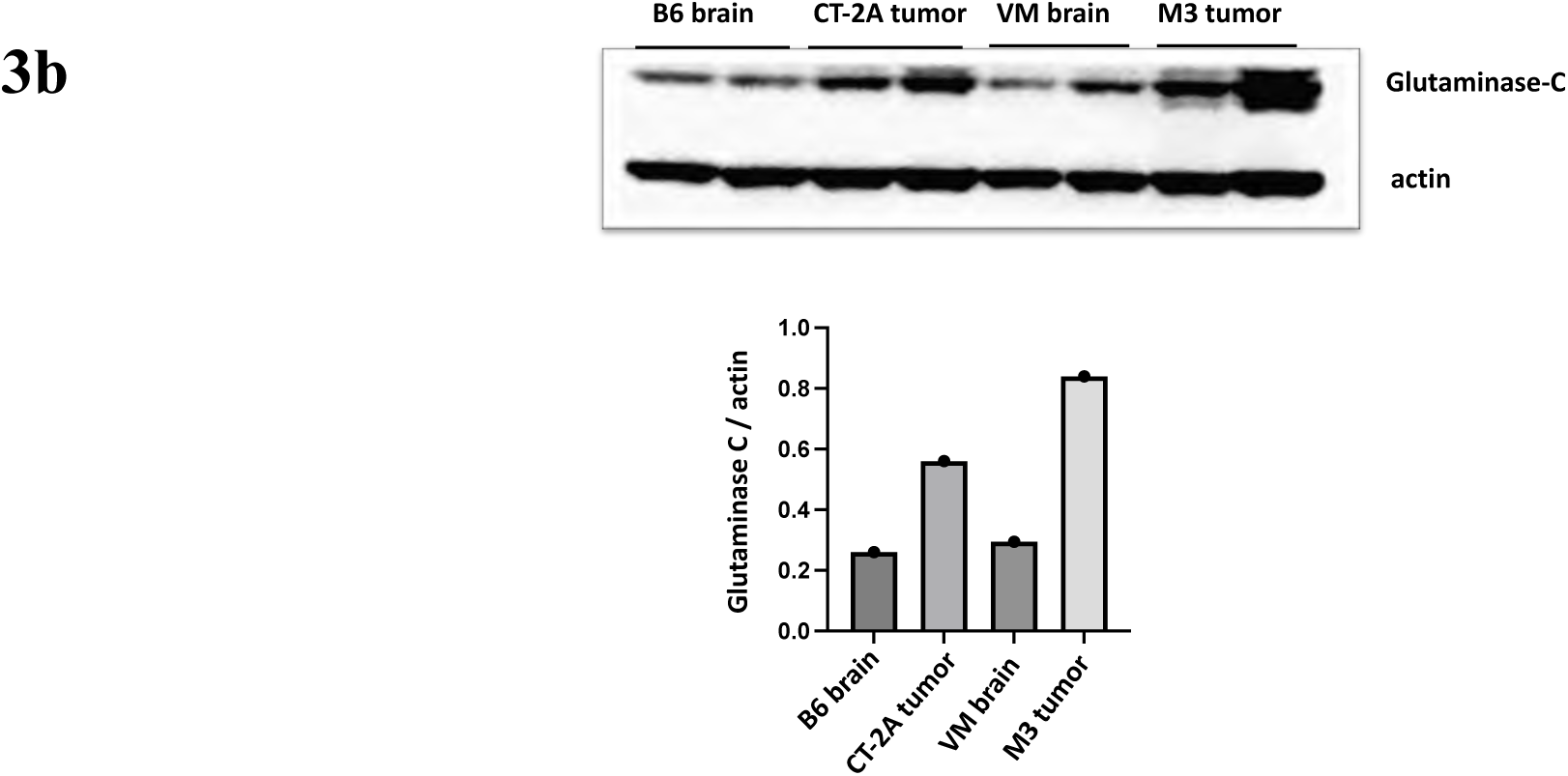
Glutaminase-C expression in normal brain and brain tumors –. Western blot analysis of glutaminase-C protein in the tissue lysates of VM-M3 and CT-2A brain tumors compared to that of corresponding normal juvenile VM and B6 brains. A markedly increase in glutaminase expression in both tumors compared to normal brains. 2 independent mice brain tissue were analyzed.

**Extended figure 4a.**
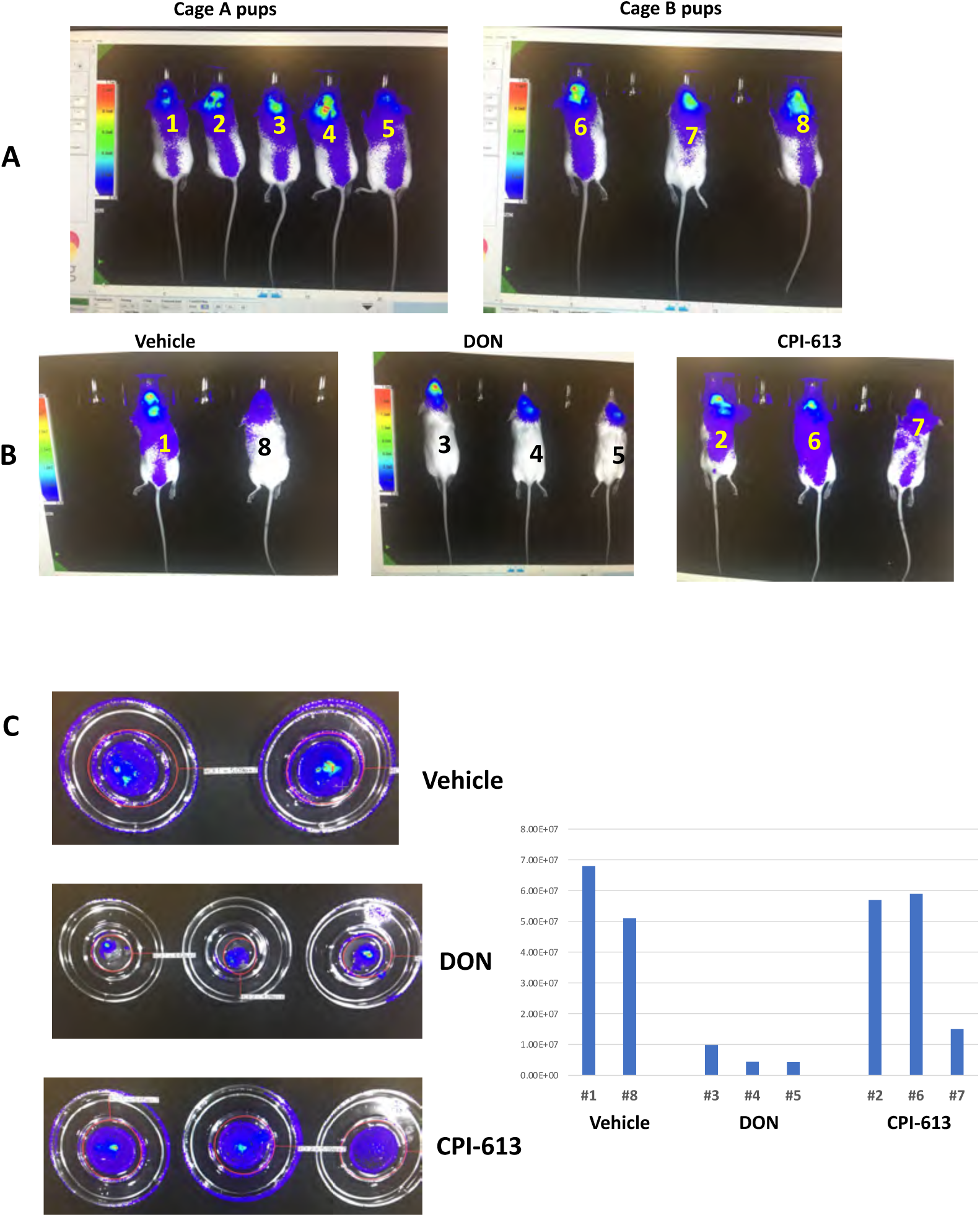
CPI-613 reduces the growth of VM-M3/luc cells *in vivo /ex vivo – a preliminary study*. VM/M3 cells were orthotopically implanted in p23 VM mice (n=8) from 2 litters (Cage A and Cage B). **Upper panel A**, mice were imaged *in vivo* 7 days after tumor implantation, numbered, and then randomly divided into 3 experimental groups: DMSO vehicle, DON (used as a positive control), and CPI-613. All mice were on an SD control diet, and drug administration started at this point as described. **Lower panel B**, mice were imaged again after day 14. **Lowest panel C**, brains were excised, imaged *ex vivo*, and quantified for bioluminescence. There were no significant differences in bioluminescence value between vehicle control and CPI-613 when the mice are under control SD.

**Extended figure 4b.**
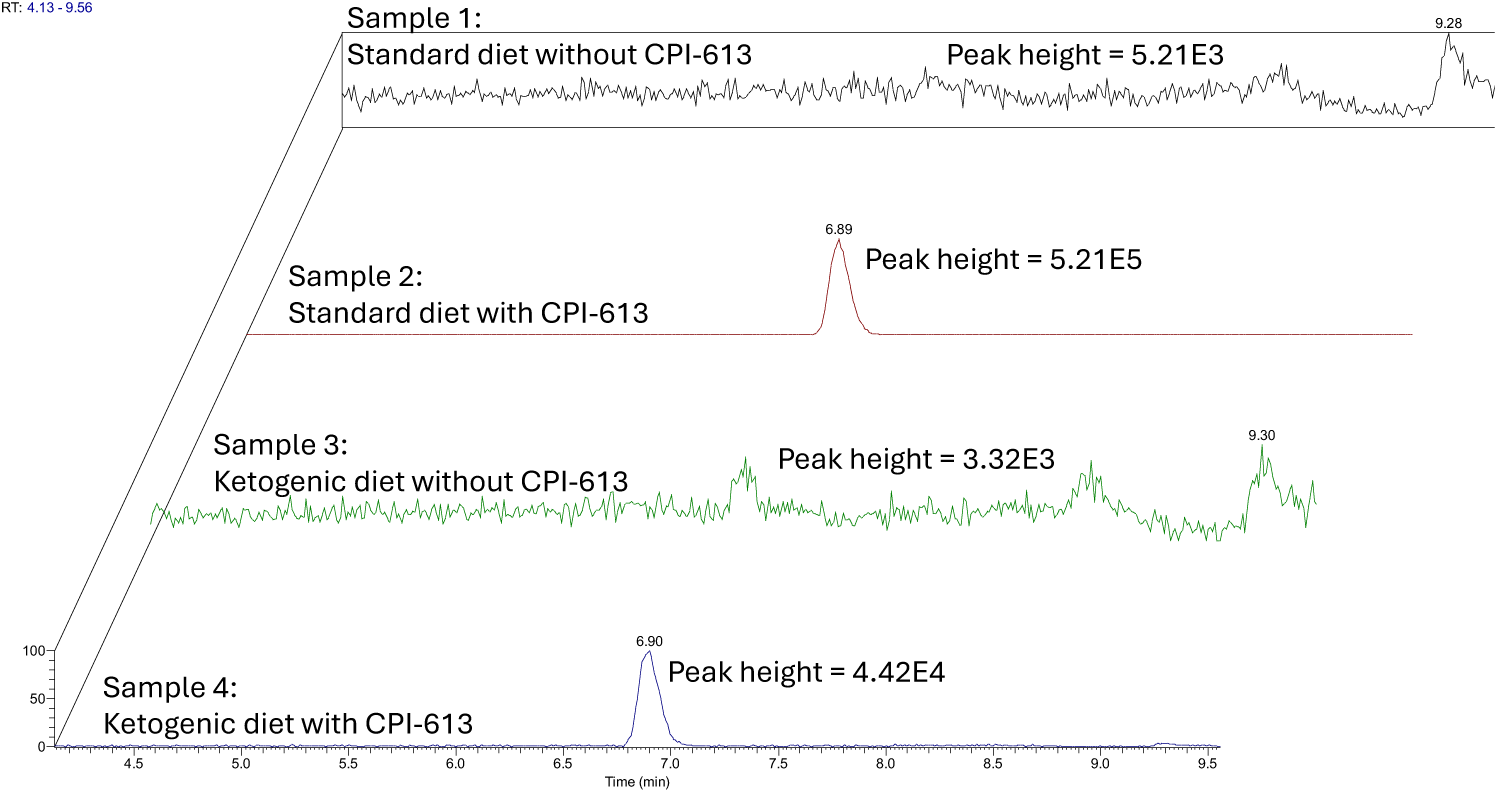
The LC-MS metabolite data of the serum showed that CPI-613 is present in both SD and KD mice. Serum was collected 45 minutes after CPI-613 injection (1mg/kg). 2 independent mice serum were analyzed.

**Extended Figure 4c.**
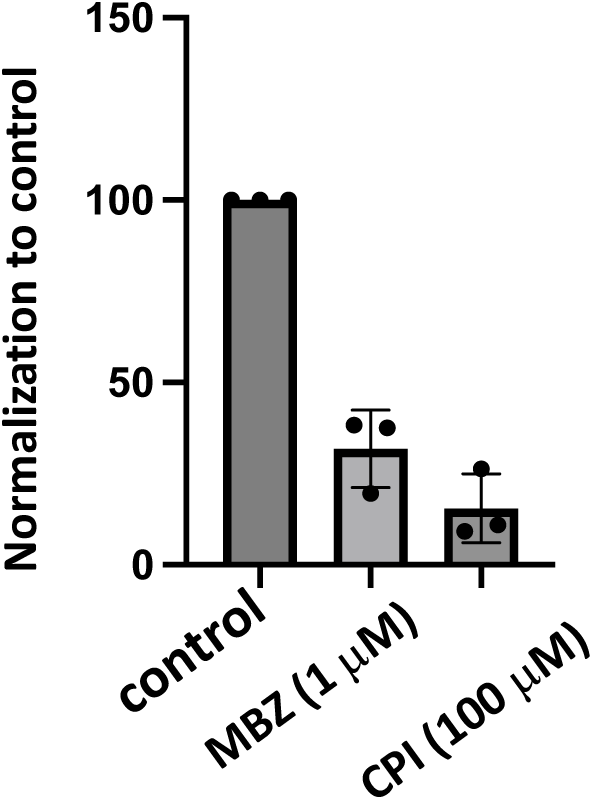
Oxygen Consumption rate (OCR) in VM-M3 cells treated with MBZ and CPI-613. VM-M3/luc cells were treated with MBZ and CPI-613 while grown *in vitro* in DMEM without serum for 6 hours as stated in results. All values are normalized and compared to control in three independent experiments. Values are plotted as mean ± SEM. OCR is significantly reduced in drug treated cells compared to control untreated cells.

**Table 1.**
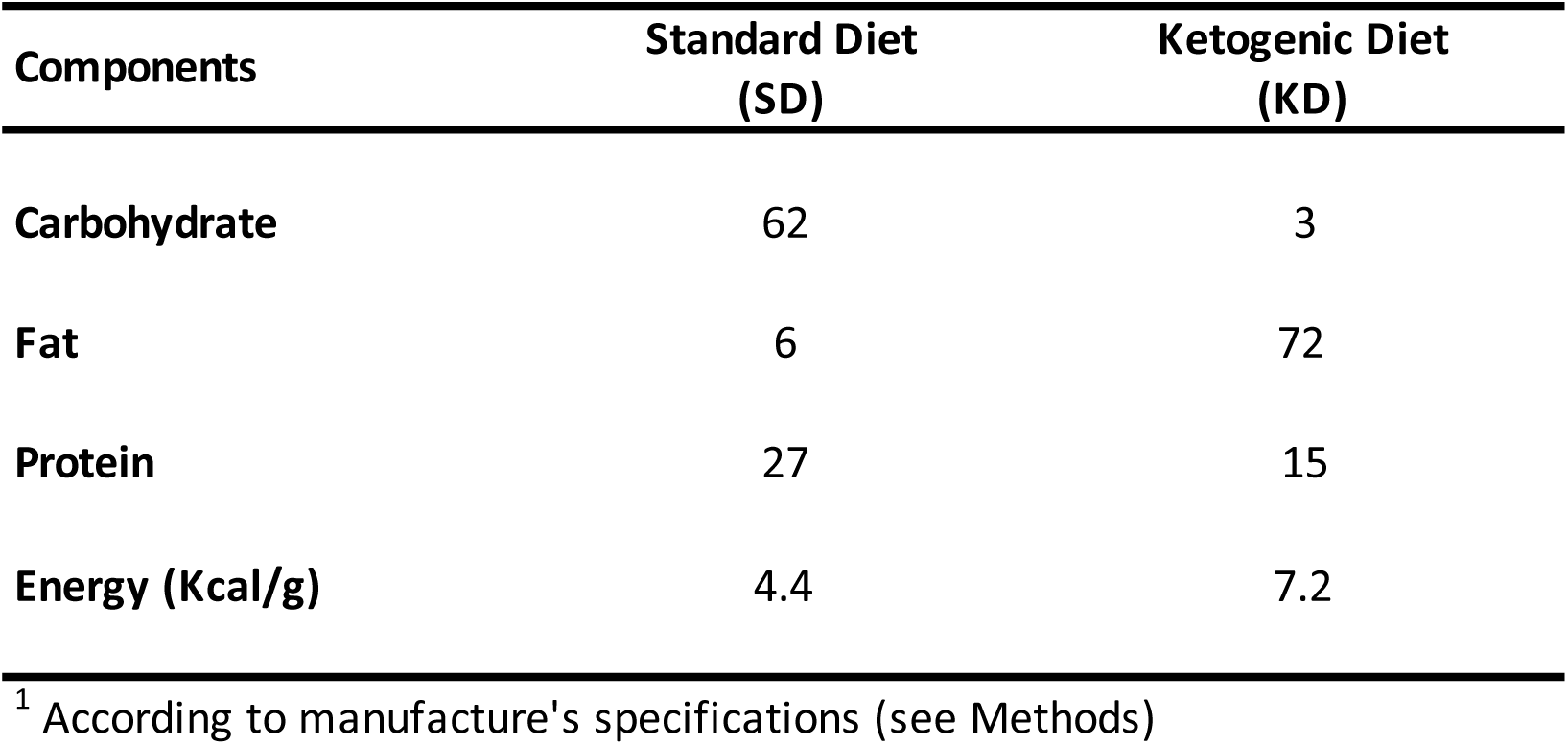
Composition (%) of the Standard diet and the Ketogenic diet^1^.

| **Components** | **Standard Diet**<br>**(SD)** | **Ketogenic Diet**<br>**(KD)** |
| --- | --- | --- |
| **Carbohydrate** | 62 | 3 |
| **Fat** | 6 | 72 |
| **Protein** | 27 | 15 |
| **Energy (Kcal/g)** | 4.4 | 7.2 |

<sup>1</sup> According to manufacture's specifications (see Methods)

## Notes

### Competing Interest Statement

The authors have declared no competing interest.

### Summary of Updates

New data, new extended figures and updated text.

