## Supplementary File 1 for "Ketogenic diet as a metabolic vehicle for enhancing the therapeutic efficacy of mebendazole and devimistat in preclinical high-grade gliomas grown in juvenile mice"

### Slide 1
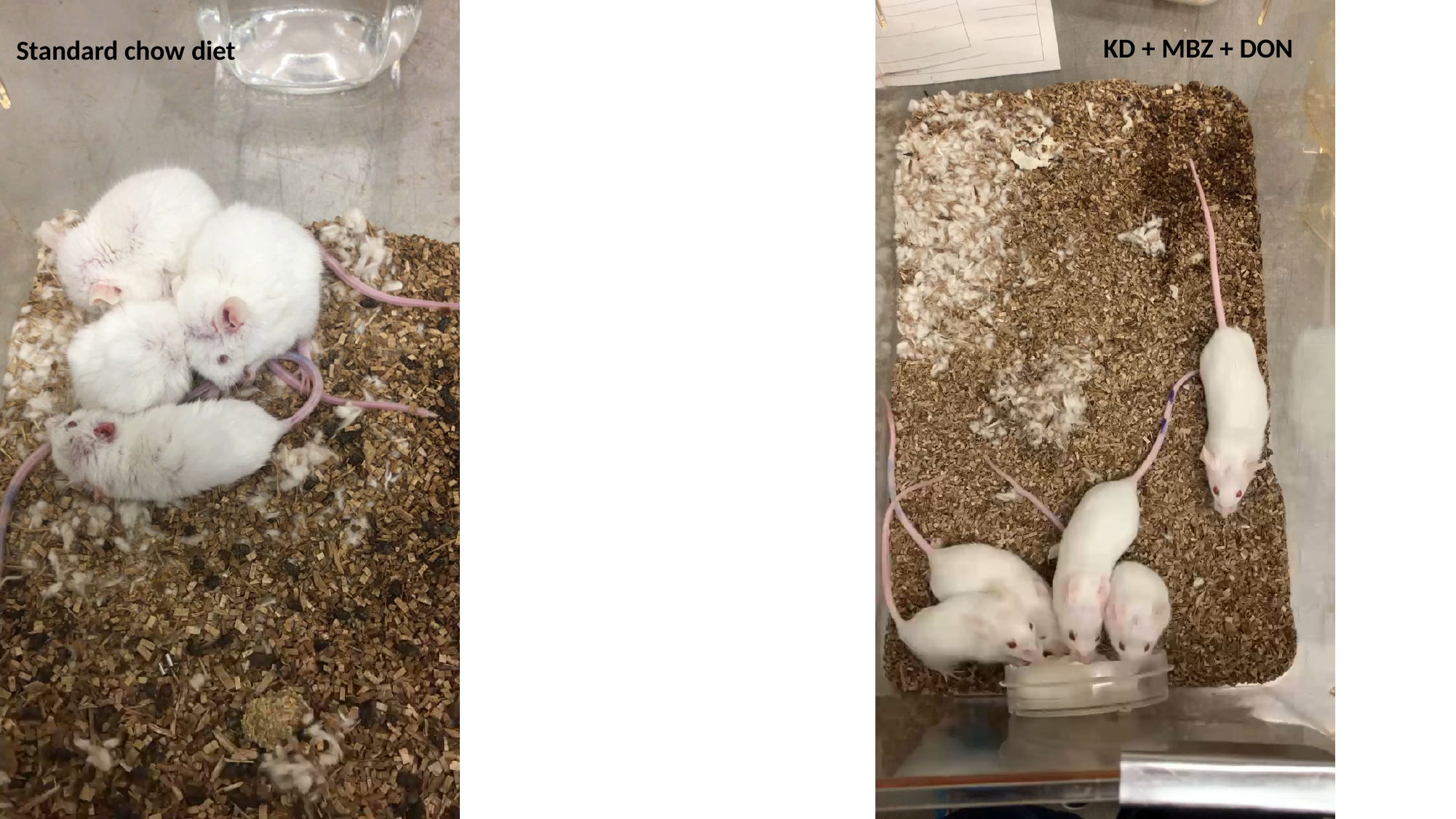

KD + MBZ + DON
Standard chow diet

### Slide 2
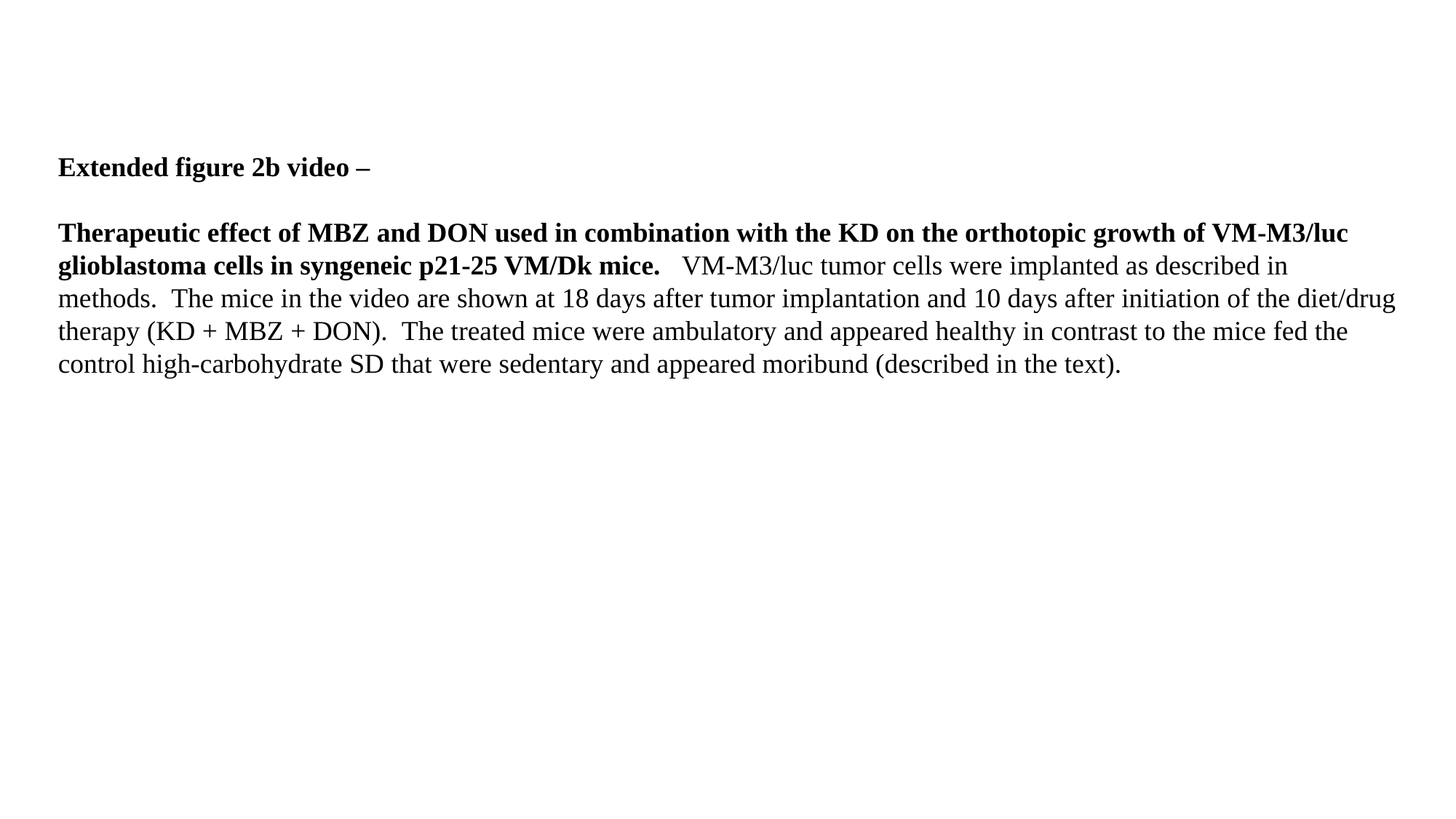

Extended figure 2b video –
Therapeutic effect of MBZ and DON used in combination with the KD on the orthotopic growth of VM-M3/luc glioblastoma cells in syngeneic p21-25 VM/Dk mice.   VM-M3/luc tumor cells were implanted as described in methods.  The mice in the video are shown at 18 days after tumor implantation and 10 days after initiation of the diet/drug therapy (KD + MBZ + DON).  The treated mice were ambulatory and appeared healthy in contrast to the mice fed the control high-carbohydrate SD that were sedentary and appeared moribund (described in the text).
